# Nuclear hormone receptor NHR-49 shapes immuno-metabolic response of *Caenorhabditis elegans to Enterococcus faecalis* infection

**DOI:** 10.1101/549907

**Authors:** Madhumanti Dasgupta, Meghana Shashikanth, Nagagireesh Bojanala, Anjali Gupta, Salil Javed, Varsha Singh

## Abstract

Immune responses to pathogenic microbes include activation of resistance and tolerance mechanisms in the host both of which are energetically expensive. In this study, we show that *C. elegans* exposed to Gram positive bacteria *Enterococcus faecalis* and *Staphylococcus aureus*, rapidly utilizes lipid droplets, the major energy reserve in the nematode. Feeding on *E. faecalis* causes developmental arrest in *C. elegans* larvae and growth arrest in adults, pointing to starvation response. We find that nematode’s early response to infection entails upregulation of 25 genes involved in lipid hydrolysis and downregulation of 13 lipid synthesis genes as early as 8 hours following exposure. We also show that lipid droplets play a protective role in *C. elegans* during infection. NHR-49, a PPARα ortholog, is required for *E. faecalis* induced beta-oxidation of fatty acids and immune effector production. It regulates an immunometabolic axis required for survival of the nematode on *E. faecalis.* Our findings reveal a facet of nutritional immunity wherein lipid droplet homeostasis plays a central role in nematode microbe interactions.

## INTRODUCTION

Host response to microbial infection requires rapid rewiring of transcriptional programs to enable production of antimicrobials providing resistance, and for activation of repair machinery providing tolerance. Both resistance and tolerance mechanisms require reallocation of resources (Rauw, 2012; Vale, Fenton and Brown, 2014). Constitutive and inducible innate defences are present in all multicellular animals. The latter entails systemic inflammatory response composed of increased production of acute phase proteins or equivalent effectors, changes in energy and nutrient metabolism, anorexia and fever in mammals (Klassing and Leschinsky, 1999; Hart, 1988) and has high energy cost. Fast living species with high reproductive rates and lower life span rely heavily on innate inducible defences and incur high cost forcing a trade off with other life history traits such as reproduction (Lee, 2006). In *Caenorhabditis elegans*, brood size is inversely correlated with immune response to pathogenic bacteria (Tang et al., 2005) suggesting that inducible defences may pose high cost in this nematode as well.

*C. elegans* in its natural habitat is exposed to a variety of bacteria, many of which are pathogenic. Immune response to pathogenic microbes entails accelerated production of large number of immune effectors such as lectins, antibacterial factors, lysozymes, saposins, caenacins and neuropeptide like proteins (Troemel et al., 2006; Irazoqui et al., 2010; Engelmann et al., 2011). The defence response is mounted as early as 4 hours and continues for at least 24 hours (Troemel et al., 2006). Even within a short time span of 4 hours, there is a dramatic increase in immune effector expression. As many as 30 immune effectors with antimicrobial or xenobiotic detoxification function are induced from 4 to 100 fold when *C. elegans* is infected by *Staphylococcus aureus* (Irazoqui et al., 2010). Rapid induction of immune effectors likely poses an increased demand on energy reserves.

Lipid droplets (LDs) are membrane enclosed organelles for storing energy in the form of neural lipids such as triacylglycerol (TAG) and cholesterol esters. During fasting, neutral LDs are broken down by lipases to release free fatty acids, which then undergo beta-oxidation to generate acetyl CoA. Acetyl CoA is utilized in tricarboxylic acid (TCA) cycle and in glyoxylate shunt (present in plants, nematodes, bacteria and fungi) to generate energy precursors and/or be utilized in gluconeogenesis. Utilization of neutral lipids is a highly orchestrated process, controlled by a complex network of conserved signalling pathways which involves many metabolic sensors and transcription factors (Leone, Weinheimer and Kelly, 1999; Van Gilst et al., 2005; Van Gilst, Hadjivassiliou and Yamamoto, 2005; Taubert et al., 2006; Nomura et al., 2010; Palanker, L., Tennessen, J.M. and Thummel, C. S., 2009). In *C. elegans*, LDs are utilized rapidly during nutrient, oxidative and cold stress (Rourke, Soukas and Carr, 2010; O’Rourke and Ruvkun, 2013; Lynn et al., 2015; Liu et al., 2017). It is unclear if LDs get utilized during infection and other biotic stresses to meet large energy demands for rapid synthesis of immune effector molecules.

In this study, we asked if *C. elegans* lipid metabolism regulates immune response to pathogenic bacteria, to impact the survival of nematodes. We find that a diet composed of Gram positive, pathogenic bacteria, *E. faecalis* or *S. aureus*, causes rapid utilization of LDs stores in *C. elegans* intestine. We show that *E. faecalis* diet causes growth arrest in *C. elegans* adults and developmental arrest in *C. elegans* larvae. We demonstrate that *E. faecalis* diet activates entire program of neutral lipid hydrolysis while suppressing neutral lipid synthesis. We also show that beta oxidation of fatty acids as well as immune effector production is dependent on a nuclear hormone receptor and PPARα ortholog, NHR-49. Increase in LD boosts survival of *C. elegans* on *E. faecalis* in an NHR-49 dependent manner. Thus, we show that survival of *C. elegans* from *E. faecalis* requires activation of NHR-49 dependent immuno-metabolic axis.

## RESULTS

### *Enterococcus faecalis* and *Staphylococcus aureus* diets induce LD depletion in *C. elegans* intestine

Nutrient stress or starvation leads to utilization of energy stored in LDs for cellular maintenance which is easily observed as decline in lipid staining in *C. elegans* intestine. To understand if there is a link between lipid metabolism and immunity, we tested if biotic stress due to feeding on pathogenic bacteria alters lipid stores in *C. elegans*. We allowed *C. elegans* adults to feed for 8 hours on two pathogenic Gram negative bacilli, *Pseudomonas aeruginosa* (PA14) and *Salmonella typhimurium* (SL1344) and on two pathogenic Gram positive cocci, *Staphylococcus aureus* (NCTC8325) and *Enterococcus faecalis* (OG1RF). *C. elegans* fed *P. aeruginosa* showed non-significant decline in lipid droplet staining, while *C. elegans* fed *S. typhimurium* showed no change in LD stores (Figure 1A and 1B). However, animals fed on *S. aureus* or *E. faecalis* (OG1RF) for 8 hours showed near complete depletion of ORO stainable lipids - similar to starvation for 8 hours (Figure 1A and 1B). Depletion of neutral lipid was also observed in 8 hours OG1RF fed *C. elegans* using neutral lipid stain BODIPY^493/503^(Figure S1A and S1B). A previous study has shown that *C.elegans* mobilize lipids to their germline during nutrient stress and oxidative stress (Lynn et al., 2015). To determine if germline plays any role in *E. faecalis* induced lipid droplet depletion, we took two approaches. In the first, we studied effect of OG1RF diet on lipid stores in very young L4 where gametocyte development and proliferation had not yet begun. We found that lipid stores were depleted in young L4 larvae on OG1RF diet (Figure S1C and S1D). In the second approach, we prevented germline proliferation by *cdc25.1* RNA interference and found that OG1RF diet caused depletion of lipid stores in these animals too (Figure 1C). Taken together, these experiments indicate that OG1RF (and *S. aureus*) feeding induces LD depletion in *C. elegans* intestine. We found that heat killed OG1RF can also induce LD depletion in *C. elegans*, as does OG1RF feeding at 20°C (Figure 1D and 1E). In all the above experiments, OG1RF was grown on nutrient rich BHI agar medium. To test for effect of bacterial growth medium, we exposed *C. elegans* to OG1RF grown on NGM agar and we observed LD depletion as well (1F). In all, our results show that OG1RF feeding induces LD depletion in *C. elegans*, in a temperature and germline independent manner.

**Figure 1.**
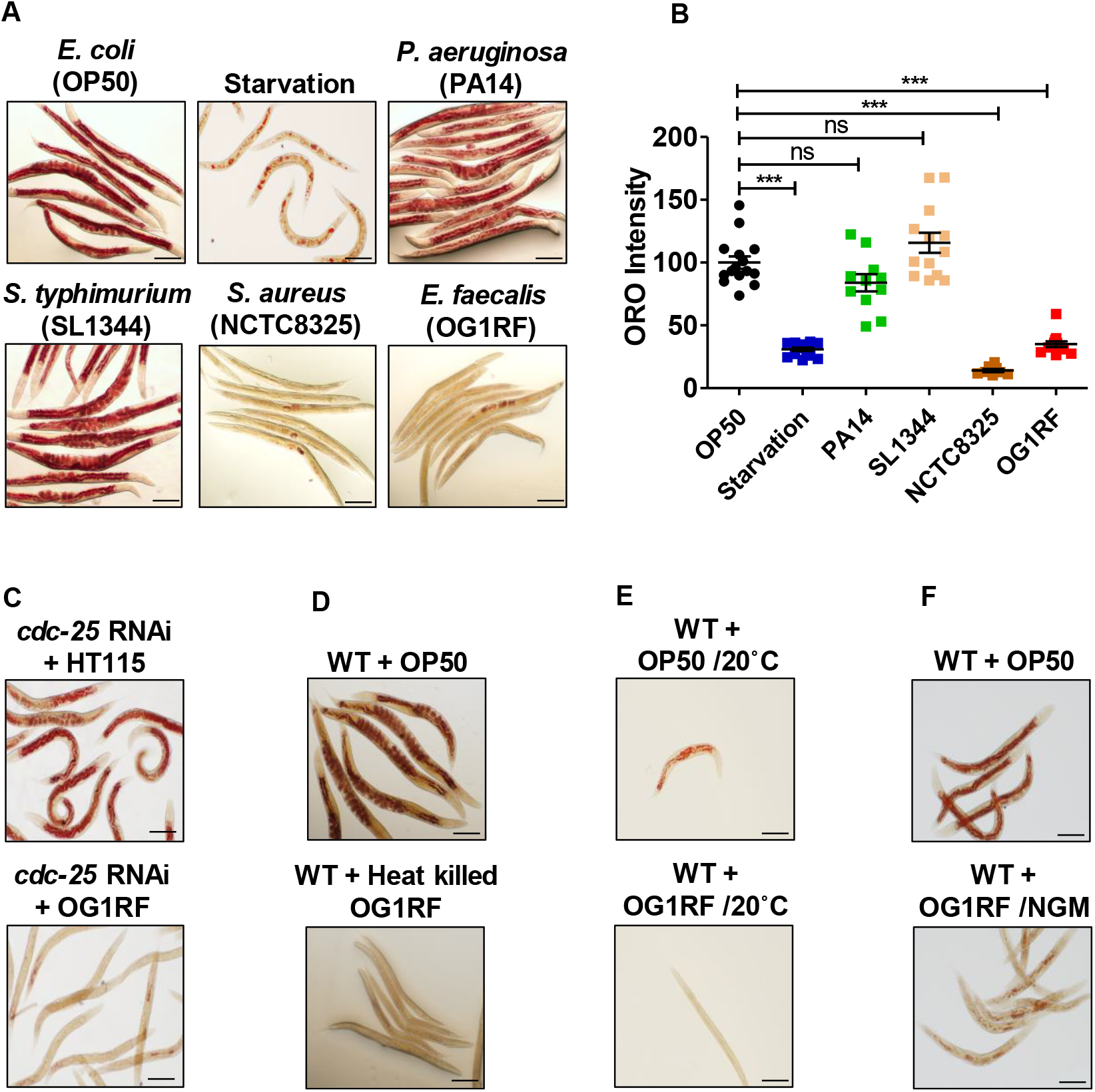
Gram positive cocci *E. faecalis* induces lipid droplet depletion in *C. elegans.* (A-B) ORO staining and quantification of adult *C. elegans* fed on *E. coli* (OP50), *P. aeruginosa* (PA14), *S. typhimurium* (SL1344), *S. aureus* (NCTC8325) and *E. faecalis* (OG1RF), or starved for 8h. (C) ORO staining of *cdc-25.1* RNAi animals fed on *E. faecalis* (OG1RF) for 8h. (D) ORO staining of animals fed on heat killed OG1RF for 8h. (E) ORO staining of animals fed on OG1RF at 20°C for 8h. (F) ORO staining and quantification of animals fed on OG1RF seeded on NGM agar for 8h. For experiments A to E, OG1RF was seeded on BHI agar. For experiments in A-D and F, feeding was performed at 25°C. Scale bar, 100 μm.

### *E. faecalis* induces larval development arrest in *C. elegans*

Nutrient limiting environment causes development delay in larvae (Cassada and Russell, 1975; Golden and Riddle, 1982) while complete starvation leads to L1 larval arrest or dauer formation (Baugh, 2013). We found that feeding on Gram negative bacteria, *P. aeruginosa* and *S. typhimurium*, did not affect larva development in *C. elegans* (data not shown). However. *C. elegans* fed on *E. faecalis* OG1RF diet had profound effect on larval development. We allowed a synchronized population of L1 larvae (250μm body length) to feed on laboratory diet *E. coli* OP50 or *E. faecalis* OG1RF lawn. We used body length as a measure to study larval development at 24, 48 and 72 hours of feeding on both bacteria (Figure 2A and 2B). We observed that *C. elegans* fed on OG1RF diet were shorter that OP50 fed animals at all time points (Figure 2B). At 72 hours of feeding, OP50 fed animals had reached body lengths measuring 1000 μm corresponding to adults while OG1RF fed animals had body lengths ranging between 250-350 μm equivalent to L1 and L2 larval stages. C

**Figure 2.**
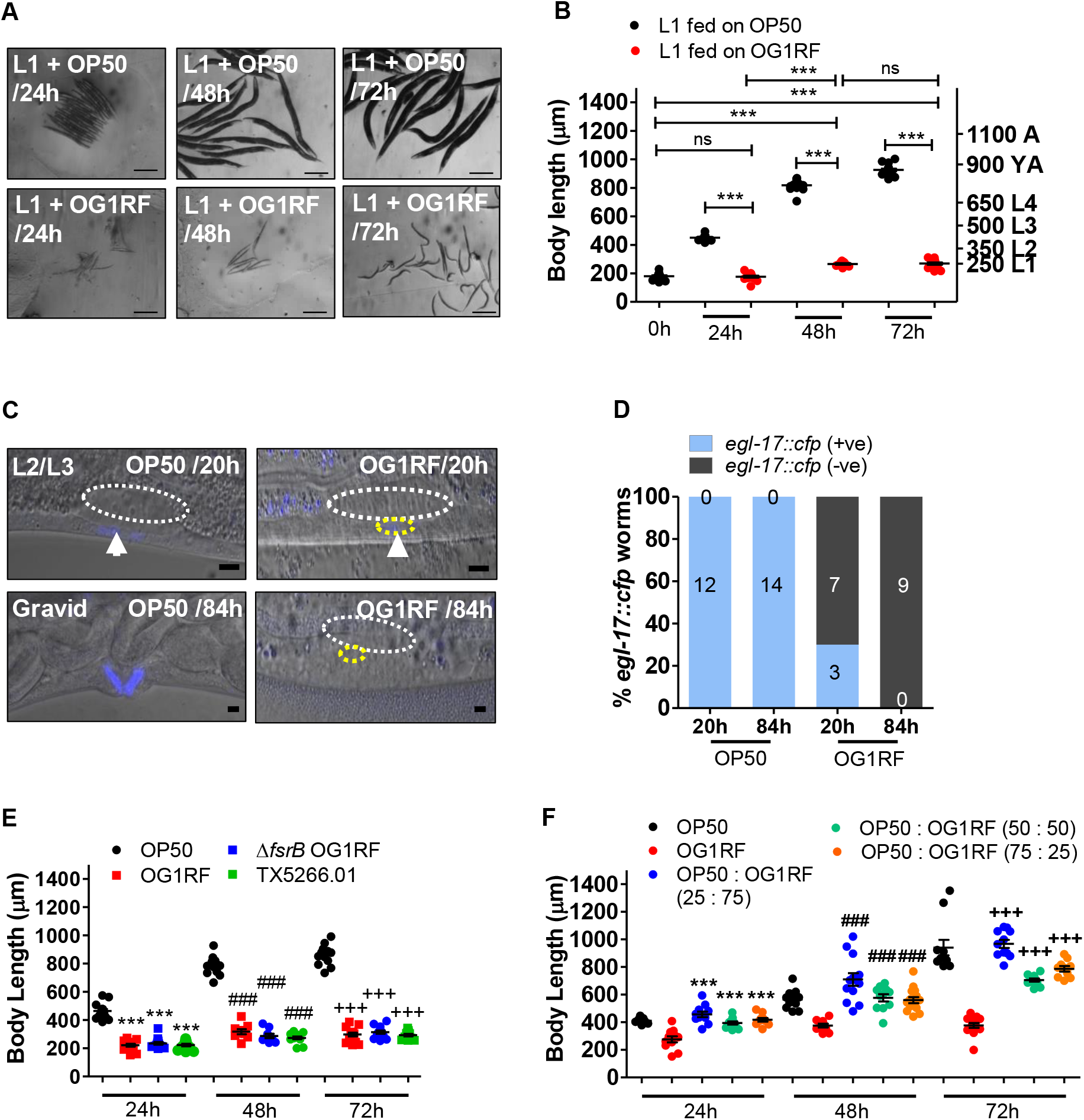
*E. faecalis* induces L2 arrest in *C. elegans.* (A) Images of *C. elegans* L1 larvae fed *E. coli* OP50 or *E. faecalis* OG1RF for indicated time. Scale bar, 100 µm. (B) Mean ± SEM of body length in larvae fed OP50 or OG1RF. (C-D) EGL-17 is targeted by inductive and notch signaling in vulva cells (Figure S2). Left panels shown are synchronized L1 larvae fed on OP50 expressing *cfp* in blue in P6.p (1°) during early L3 (20h) and in VulD (2°) in gravid animals (84h). OG1RF fed larvae show weak expression of *egl*-17::cfp at 20h (white arrow head) and no expression in undivided P6.p (white arrow) 84h. White circles depict gonad primordium and yellow P6.p. In all animals anterior is to the left. Scale bar, 5 µm. (D) Quantification of *egl-17*p::cfp in OP50 and OG1RF fed animals at 20h and 84h. (E) Mean ± SEM of body length in L1 larva fed on OP50, OG1RF, *ΔfsrB* OG1RF (TX5266) and the complemented strain TX5266.01. (F) Mean ± SEM of body length of L1 larvae fed on mixed lawn of OP50 an OG1RF.

To confirm OG1RF induced arrest of larval development, we studied proliferation and lineage progression in two organs, hermaphrodite gonad and vulva. Development of both these organs is tightly synchronized with larval development. In *C. elegans* hermaphrodite, vulva development from vulval precursor cells (VPC) is dependent on signalling from the gonadal cells called Anchor Cell (AC). AC secretes LIN-3/EGF (Epidermal Growth Factor) to induce vulva fate in VPCs- P5.p, P6.p and P7.p-of the equipotent P(3-6).p lineage cells. P6.p, the VPC closest to AC adopts 1^0^cell fate, and VPCs P5.p/P7.p adopt 2^0^fates. The EGL-17/FGF (Fibroblast Growth Factor) is induced transcriptionally by inductive and notch signalling in vulva cells (Sulston and Horvitz, 1977; Burdine, Branda and Stern, 1998) From early L3 to late L3, egl-17 is expressed in P6.p/1^0^lineages and at mid L4 it switches to 2^0^lineage cells, VulC and VulD (Figure S2A and S2B). We followed P6p fate in OP50 and OG1RF fed larvae. At 20h post feeding (time prior to VPC P6.p division), 100% of OP50 fed larvae showed positive *egl-17*::*cfp* expression corresponding to L3 larval stage. However, in OG1RF fed larvae only 30% larvae showed weak expression in P6.p cell corresponding to late L2 stage while 70% had no GFP expression suggesting an early L2 stage. At 84 hrs of feeding, we observed *egl-17*::*cfp* expression in vulD corresponding to adult stage in OP50 fed animals, while no cell division in P6.p lineage was observed in OG1RF fed larvae (Figure 2C and 2D). These observations confirm that the vulva development in OG1RF fed larvae was indeed arrested at late L2 as VPCs did not divide and never acquired vulva fates.

Gonadogenesis in *C. elegans* hermaphrodite starts with four cells in the primordium or gonad pouch at the time of hatching of L1. The primordium proliferates till adulthood via somatic and germline proliferations, and stereotypic migrations of gonadal arms guided by Distal Tip Cells (DTCs) (Kimble and Hirsh, 1979) (Figure S3A and S3B). To follow gonad development in OG1RF fed larvae, we counted nuclei in gonad primordia with DIC microscopy. Control, OP50 fed, larvae showed 12 nuclei at 12 hours (mid L2), complete gonad development at 36 hours (late L4) and at 60 hours (gravid adult) (Figure S3C-D). But, OG1RF fed larvae showed either 4 nuclei (L1) and 12 or more nuclei (L2) at 12h hours (L1=4/17, L2=13/17) as well as at 36 hours (L1=15/23, L2=17/23). OG1RF fed larvae had only L2 gonad primordia even at 60 hours (n=20) time point (Figure S3C-D). This indicated that gonad development was also arrested at L2 stage in OG1RF fed larvae of *C. elegans*. Also, we could not find larvae with L3 gonad features such as germline proliferation or extension or bending of gonad arms in any of the OG1RF fed larvae. In all, three different parameters of growth and development-body length measurements, vulva development and gonad development-demonstrate that OG1RF induces L2 arrest in *C. elegans*.

We set out to ask if larval arrest on OG1RF diet was due to nutrient restriction/starvation or a toxin mediated effect. To test if *E. faecalis* pathogenesis was a cause for arrest, we utilized a virulence attenuated strain of OG1RF, *ΔfsrB* OG1RF and *fsr* complemented OG1RF strain in larval development assay (Qin et al., 2000; Garsin et al., 2001). Both strains caused larval arrest similar to the arrest by OG1RF feeding, suggesting that pathogenesis is not the only cause of arrest (Figure 2E). If poor nutrition from OG1RF diet was responsible for arrest, supplementation of OG1RF diet with OP50 cells should alleviate larval arrest. We allowed L1 larvae to feed on monoxenic lawns of OP50 and OG1RF or on mixed diet of OP50 and OG1RF. We found that replacing as little as 25% of OG1RF cells with OP50 cells could suppress the larval arrest induced by OG1RF alone (Figure 2F).

To test if OG1RF induced larval arrest was reversible, we allowed 48h starved larvae, 48h OG1RF fed larvae and 48h OP50 fed larvae to resume/continue feed on OP50 for further 12h, 24h, 36h, 48h and 72h (Figure S4A and S4B). As shown, both starved and OG1RF arrested larvae resumed larval development when fed OP50 diet, although starved larvae were slower in resuming development. Taken together, body length measurement, gonad development and vulva development assay, we show that OG1RF diet induces larval arrest in *C. elegans*, at the L2 stage. The arrest was reversible and could be rescued by providing OP50 cells as supplement.

### *E. faecalis* diet leads to growth delay in adult *C. elegans*

Reversible and rescuable nature of OG1RF induced developmental larval arrest in *C. elegans* suggested that poor nutrition may contribute to arrest. Poor nutrition should also affect growth in adult *C. elegans*. To test this, we allowed L4s to feed on OP50 or OG1RF for 8 and 24 hours. L4s displayed an average increase of 130 µm in 8 hours and 250 μm in 24 hours when fed on OP50, but there was no significant increase in body length in 8 hours and a mere increase of 90 μm in 24 hours when fed on OG1RF (Figure 3B). We also observed OG1RF induced delayed adult growth even in *cdc-25.1* RNAi animals (Figure S4). Lack of increase in body length prompted us to examine macronutrient content in OG1RF fed animals. We measured carbohydrate content, by Anthrone test, and protein content, by Bradford test, in OG1RF and OP50 fed *C. elegans* (Figure 3C and 3D). Although protein levels remained similar in OP50 and OG1RF fed animals, we found that OG1RF fed animals had 50% lower carbohydrate content compared to OP50 fed animals (Figure 3C).

**Figure 3.**
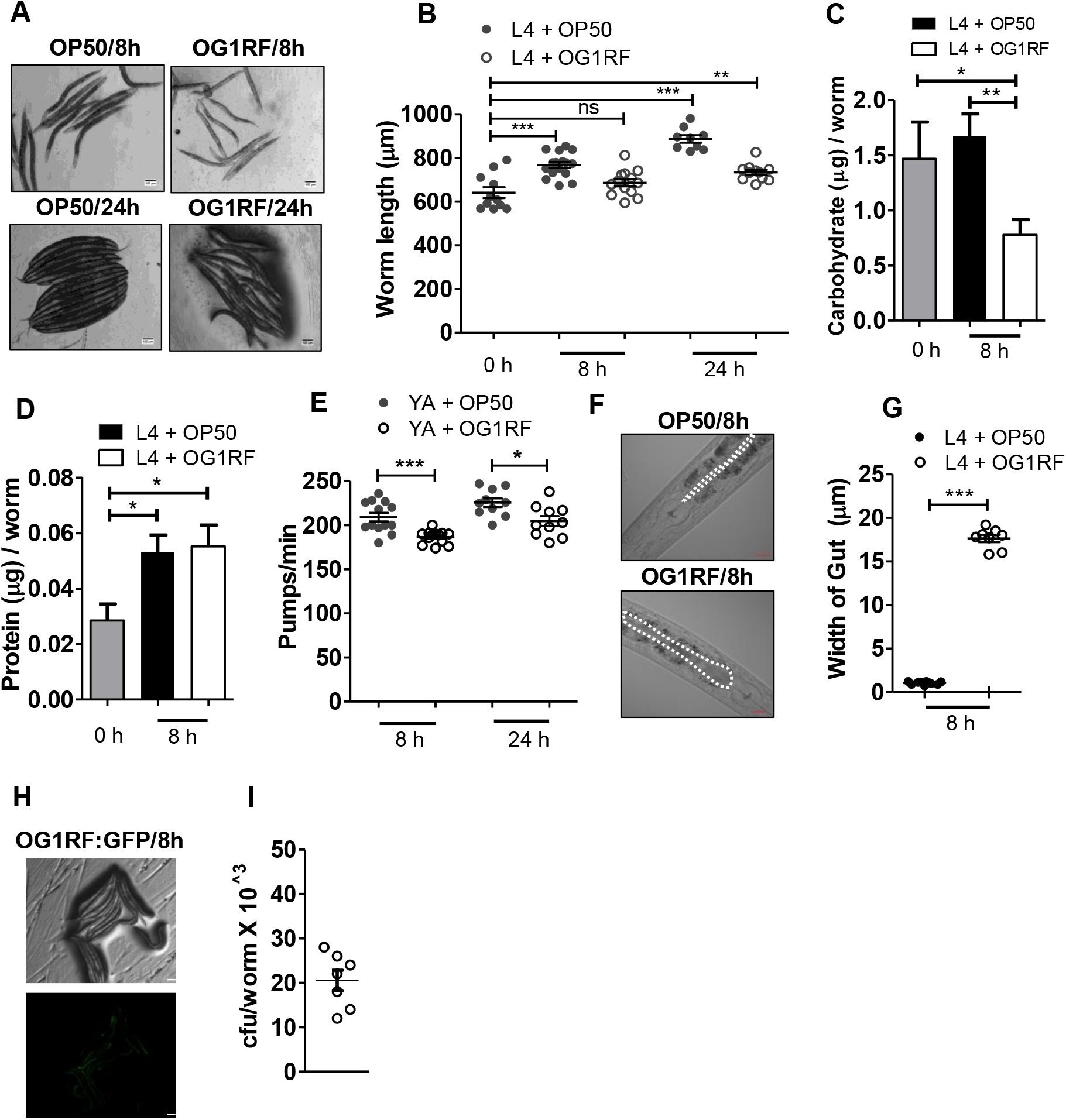
*E. faecalis* diet causes growth delay in adult *C. elegans.* (A) Images of L4 animals fed OP50 and OG1RF for 8h and 24h. Scale bar, 100μm. (B) Body length of *C. elegans* adult fed OP50 or OG1RF. Mean ± SEM of (C) Carbohydrate and (D) protein in L4 animals fed OP50 and OG1RF for 8h. (E) Pharyngeal pumps per minute in young adult worms fed OP50 or OG1RF for 8h and 24h. (F-G) DIC images of gut lumen distention in L4 animals fed on OP50 or OG1RF for 8h. (G) Mean ± SEM of gut lumen width in OP50 and OG1RF fed animals. (H-I) OG1RF:GFP colonization in *C. elegans* adult fed for 8h. Scale bar, 100μm. (I) Colony forming units of OG1RF:GFP recovered from 8h fed animals.

To account for loss in carbohydrate content in OG1RF fed animals, we tested rate of feeding. We found that there was a very small decline (10%) in pharyngeal pumping on OG1RF than on OP50 (Figure 3E). In fact, we found that *C. elegans* intestinal lumen were full of cocci suggesting that *C. elegans* was able to ingest OG1RF. Consequently, we found that intestinal lumen was distended to around 17 μm when animals were fed OG1RF, whereas no such distention was found in OP50 fed animals (Figure 3F and G). We could recover an average of 20,000 cocci per animal in 8 hours of feeding suggesting that *C. elegans* can ingest a large number of cocci (Figure 3H and I). Taken together, our results suggest that *C. elegans* ingests OG1RF, but this diet poorly supports growth pointing to possible lack of nutrition on *E. faecalis* diet.

### *E. faecalis* diet activates a transcriptional program for neutral lipid utilization in *C. elegans*

It is well documented that lipid droplet utilization during starvation of *C. elegans* requires activation of lipid hydrolysis pathways. This includes upregulation in the levels of lipases and enzymes involved in beta oxidation of fatty acids (Van Gilst, Hadjivassiliou and Yamamoto, 2005; Larance *et al.*, 2015). We hypothesized that rapid depletion of neutral lipids in OG1RF fed animals requires upregulation of transcripts for key enzymes involved in lipid breakdown. Therefore, we analysed the transcriptional response of *C. elegans* fed on OG1RF and OP50 by RNA sequencing. We identified 1750 differentially expressed genes (1130 upregulated, 619 downregulated) when animals were fed on OG1RF for 8 hours compared to OP50 (Table S2). Of the upregulated genes, 25 were involved in various steps of lipid hydrolysis (Figure 4A) and are highlighted in lipid breakdown and utilization schematic in Figure 4B (see Table S2 for fold changes).

**Figure 4.**
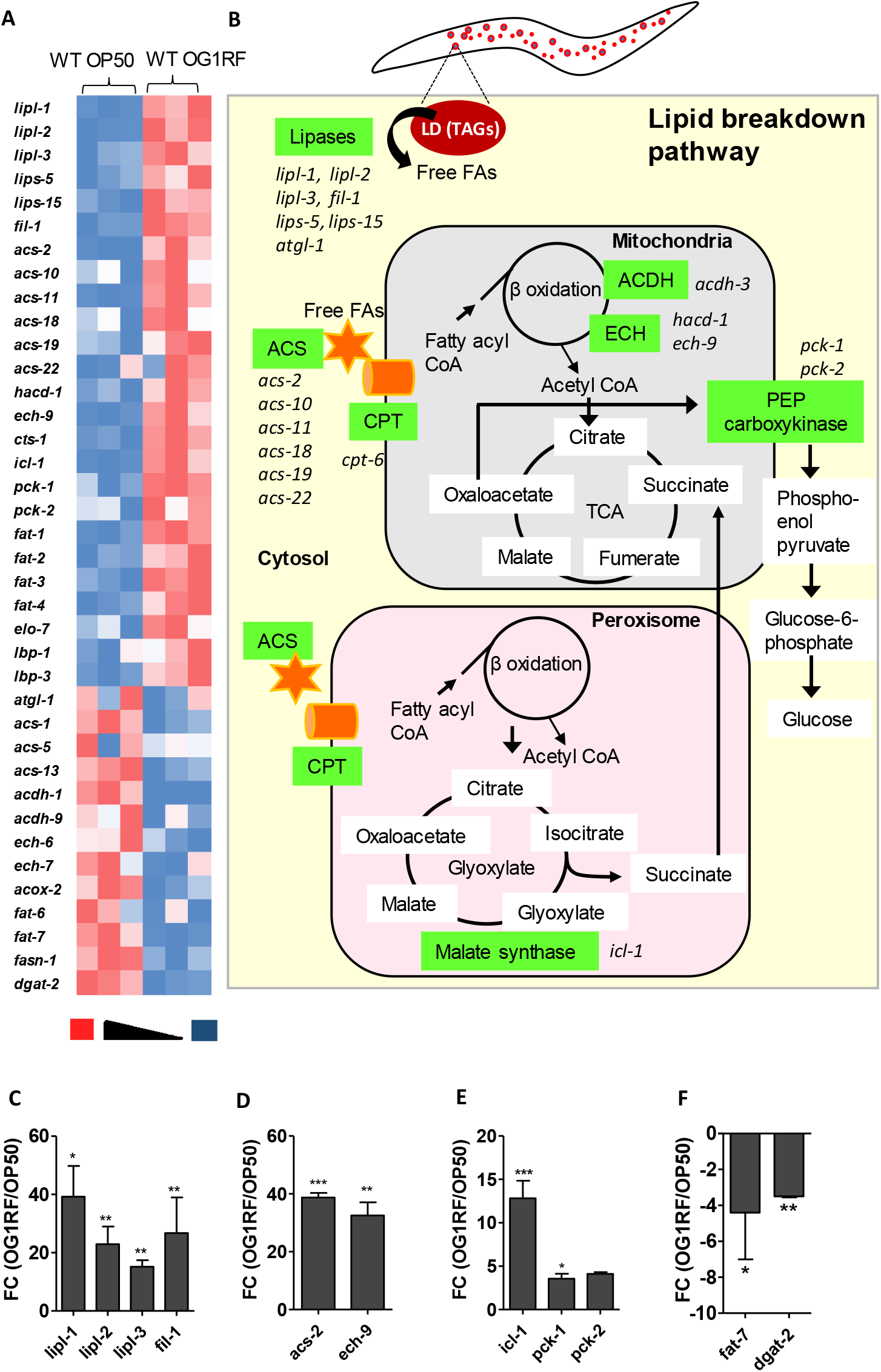
*E. faecalis* diet activates a transcriptional program for neutral lipid utilization in *C. elegans*. (A) Heat map of expression of lipid metabolism genes in *C. elegans* L4 larvae fed on OP50 and OG1RF for 8h. Red indicates upregulation and blue indicates downregulation. (B) Schematic of lipid metabolism in *C. elegans*. Genes which show upregulation in transcript level on OG1RF diet are highlighted in green boxes. Quantitative real time PCR analysis of (C) lipases, *lipl-1, lipl-2, lipl-3* and *fil-1* (D) Beta oxidation genes *acs-2* and *ech-9*, (E) *icl-1* and *pck-1*, and (F) *fat-7* and *dgat-2* in WT animals fed on OG1RF diet for 8h, compared to OP50 fed animals.

We validated RNA sequencing data using quantitative real time PCR (Figure 4C, D and E, and Figure S6). Lipid hydrolysis requires sequential action of lipases on triacylglycerides. Transcripts for four lipases, *lipl-1, lipl-2, lipl-3* and *fil-1-* were upregulated during OG1RF feeding. Free fatty acids released by lipases are charged with a coenzyme A moiety by acyl CoA synthetase enzyme (ACS family) before their transport into the mitochondrion or the peroxisome for beta oxidation. Of particular interest was acyl CoA synthetase *acs-2* which was dramatically upregulated, over 40-fold, in OG1RF fed animals compared to OP50 fed animals (Figure 4B and D). In addition, transcript for one beta oxidation enoyl-CoA hydratase *ech-9* was also upregulated 32-fold (Figure 4D). Beta oxidation of individual fatty acids in the mitochondria, or in the peroxisomes leads to production of acetyl CoA for use in TCA cycle, glyoxylate shunt or gluconeogenesis as outlined in Figure 4B. Thus, transcriptome analysis showed that OG1RF diet induces a broad transcriptional response to activate sequential steps of lipid hydrolysis in *C. elegans*. We also found three beta oxidation genes-*acdh-1, acdh-2* and *ech-6* downregulated and three desaturases required for PUFA synthesis-*fat-1, fat-2* and *fat-3* upregulated in OG1RF fed animals (Figure S5). This is a similar to pattern of dysregulation seen during starvation (Van Gilst, Hadjivassiliou and Yamamoto, 2005).

Acetyl CoA generated by fatty acid beta oxidation can be taken up for ATP production via TCA cycle in the mitochondria or the glyoxylate shunt in the peroxisome. Intermediates generated from both the pathways can move into gluconeogenesis via phospho-enol pyruvate carboxykinase (PEPCK). *G*lyoxylate shunt, in *C. elegans*, plants and bacteria, generates two molecules of ATP per acetyl CoA molecule like the TCA cycle, without the loss of molecule of CO^2^ that happens in TCA. Isocitrate lyase ICL-1/malate synthase is the rate limiting enzyme of glyoxylate shunt. We found that *icl-1* transcripts were induced >10 fold in OG1RF fed animals (Figure 4E). Transcript for rate limiting enzyme of gluconeogenesis, PEPCK encoded by *pck-1* was also upregulated ∼3 fold in OG1RF fed animals (Figure 4E). Expression of *pck-2*, another isoform of *pck-1*, was also induced in OG1RF fed animals (Table S2). Activation of lipid breakdown accompanied downregulation of Δ9 desaturase *fat-7*, and of diacyl glycerol acyl transferase, *dgat-2*, rate limiting enzyme for lipid droplet synthesis (Figure 4F). Downregulation of these genes coincide with activation of lipid breakdown (Van Gilst, Hadjivassiliou and Yamamoto, 2005). In all, the transcriptome analysis suggests that *C. elegans* fed on OG1RF for 8 hours indeed engage in a “starvation like” transcriptional response to enable them to derive energy, from neutral lipid stores in the intestine, for sustenance and possibly also for immune response.

### NHR-49 shapes metabolic response as well as immune response in *E. faecalis* fed *C. elegans*

LD homeostasis is a highly regulated process involving a number of conserved metabolic sensors and transcription factors in *C. elegans*. We tested some of the known regulators of lipid homeostasis modulate metabolic rewiring we observed in C. elegans on OG1RF diet (figure 4D). Using an *acs-2P::GFP* reporter, we observed a strong induction in reporter expression in *C. elegans* adults fed on OG1RF for 8 hours (Figure 5A and 5B), recapitulating qPCR and RNAseq data. We found that *acs-2P::GFP* reporter was induced by Gram positive bacterium, *S. aureus* but not by Gram negative bacteria *P. aeruginosa* or *S. typhimurium* (Figure S7). We tested for the requirement of transcription factors (SBP-1, NHR-80, NHR-49) and coactivator MDT-15 in OG1RF induced upregulation of *acs-2P::GFP*. We found that NHR-80 and SBP-1 were not required for reporter induction on OG1RF diet, but both NHR-49 and coactivator MDT-15 were necessary (Figure 5C).

**Figure 5.**
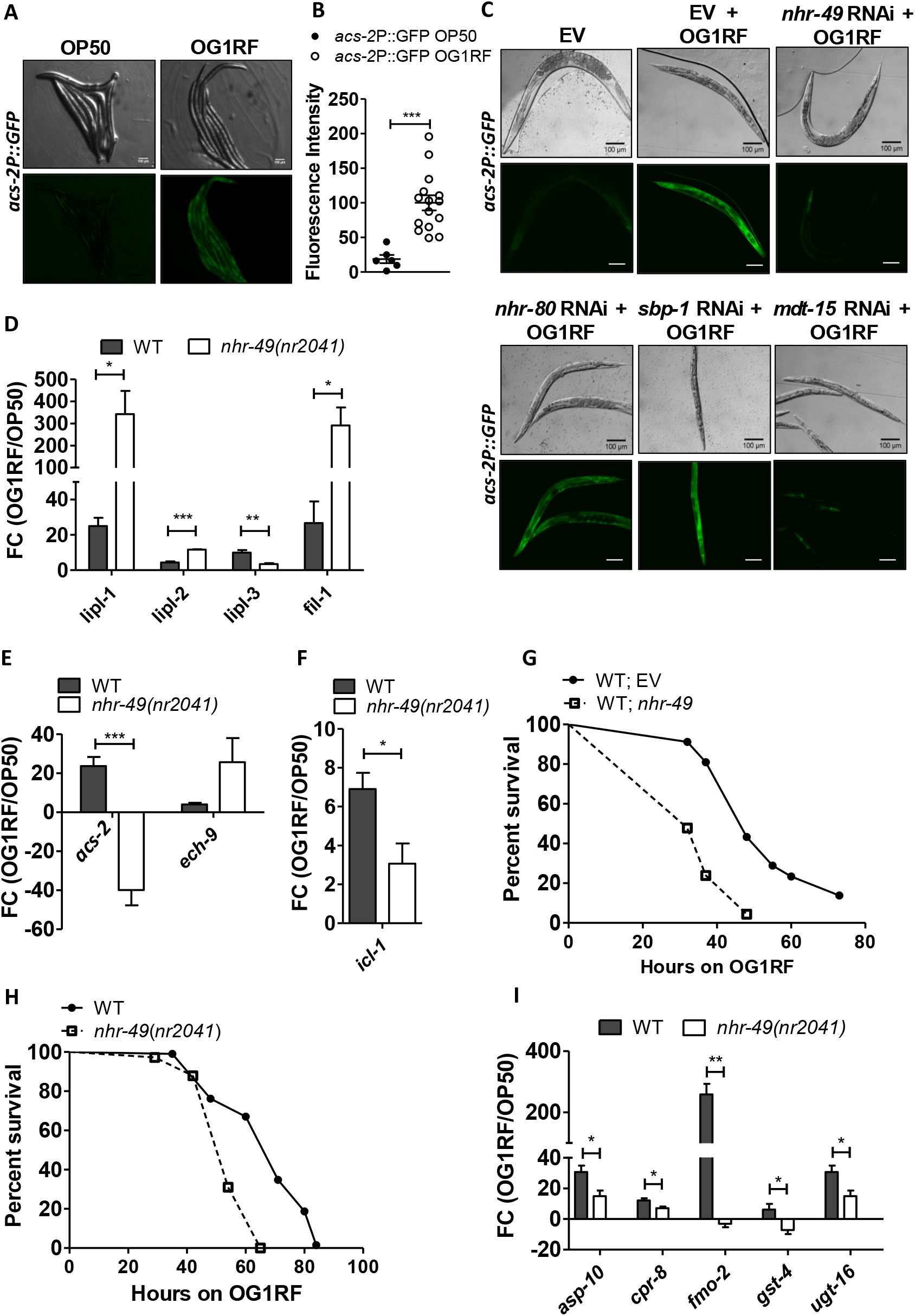
NHR-49 shapes the metabolic as well as immune response in *C. elegans* fed on *E. faecalis.* (A) Images of *acs-2P::GFP* animals fed OG1RF or OP50 for 8h. (B) Mean ± SEM values for *acs-2P*::fluorescence in OP50 and OG1RF fed animals. (C) Effect of RNAi of *nhr-80, sbp-1, mdt-15, nhr-49* on *acs-2P::GFP* fluorescence upon feeding on OG1RF for 8h. Quantitative real time PCR analysis of OG1RF induced expression of (D) lipases, (E) *acs-2* and *ech-9*, (F) *icl-1* in nhr-49(nr2041) animals compared to WT animals. (G) Kaplan Maier survival curve of (G) *nhr-49* RNAi animals and control animals exposed to OG1RF (P< 0.0001), (H) *nhr-49(nr2041)* and WT animals exposed to OG1RF (P< 0.0001). (I) Quantitative real time PCR analysis of OG1RF induced expression of immune effectors in WT and *nhr-49(nr2041)* animals fed OG1RF for 8h compared to OP50.

NHR-49 is an ortholog of human peroxisome proliferator activated receptor α (PPARα) and regulates expression of many lipid breakdown enzymes including *acs-2* and *gei-7/icl-1* (Van Gilst, Hadjivassiliou and Yamamoto, 2005; Ratnappan et al., 2014). Using an *nhr-49* deletion mutant, *nhr-49(nr2041)*, we confirmed that OG1RF induced upregulation of *acs-2* transcript relied on NHR-49 (Figure 5E). We examined whether other lipid hydrolysis genes were dysregulated in *nhr-49(nr2041)* animals. Indeed, we found that apart from *acs-2*, NHR-49 activity was required for upregulation of *lipl-3* (lipase) and *icl-1* (isocitrate lyase) on OG1RF diet (Figure 5D, E and F). Surprisingly we found that there was an induction in transcript levels of lipases, *lipl-1, lipl-2* and *fil-1*, and enoyl CoA hydratase, *ech-9* in *nhr-49*(*nr2041*) in animals on OG1RF (Figure 5D, E and F).

RNAi inhibition of *nhr-49* (Figure 5G) or mutation (Figure 5H) also made *C. elegans* adults susceptible to death induced by chronic exposure to OG1RF as shown earlier (Sim and Hibberd, 2016). We hypothesized that NHR-49 regulates immune response by facilitating lipid droplet hydrolysis for energy to promote immune effector production. Several immune effectors-antimicrobial factors, detoxification enzymes are upregulated during OG1RF infection as seen in transcriptome data (Table S2). We found that expression of five transcripts involved in immune response- UDP-glycosyltransferase (*ugt-16*), flavin mono-oxygenase (*fmo-2*), cysteine protease (*cpr-8*), aspartyl protease (*asp-10*) and glutathione-s-transferase (*gst-4*)-were dampened in *nhr-49*(*nr2041*) animals compared to wild type animals exposed to OG1RF (Figure 5F). We tested expression of additional immune effector transcripts but found no significant change in expression of cysteine protease (*cpr-4)*, cyclophilin *(cyp-35C1)* and inducible lysozyme (*ilys-2*) between wildtype and mutant animals, however saposin (*spp-2)* and F53A9.8 had high expression levels in the *nhr-49(nr2041)* animals (Figure S8). Taken together, transcriptome analysis showed that NHR-49 regulates both lipid catabolism and immune effector production in adult *C. elegans* fed on *E. faecalis*.

### Neutral Lipids boost *C. elegans* survival during *E. faecalis* infection

Neutral lipid breakdown provides energy via glyoxylate shunt, and provides carbon for gluconeogenesis, to endure a period of nutrient stress. We reasoned that neutral lipids could also be utilized to boost effector production during stress. We hypothesized that animals with increased LD content can better endure starvation and can mount a more effective immune response when fed OG1RF. We utilized a regime of 10 mM glucose supplementation to OP50 diet for only 2 days, during development, to increase lipid content in *C. elegans* intestine as shown before (Han et al., 2017). Glucose supplementation prior to infection led to 30% more ORO staining in adults than control (Figure 6A and B). Glucose supplemented adults, with higher ORO stained lipids, showed modest but significant increase in survival upon chronic exposure to OG1RF than animals with normal LD stores (Figure 6C). This indicated that increase in lipid stores had beneficial effect upon enduring pathogenesis of *E. faecalis*.

**Figure 6.**
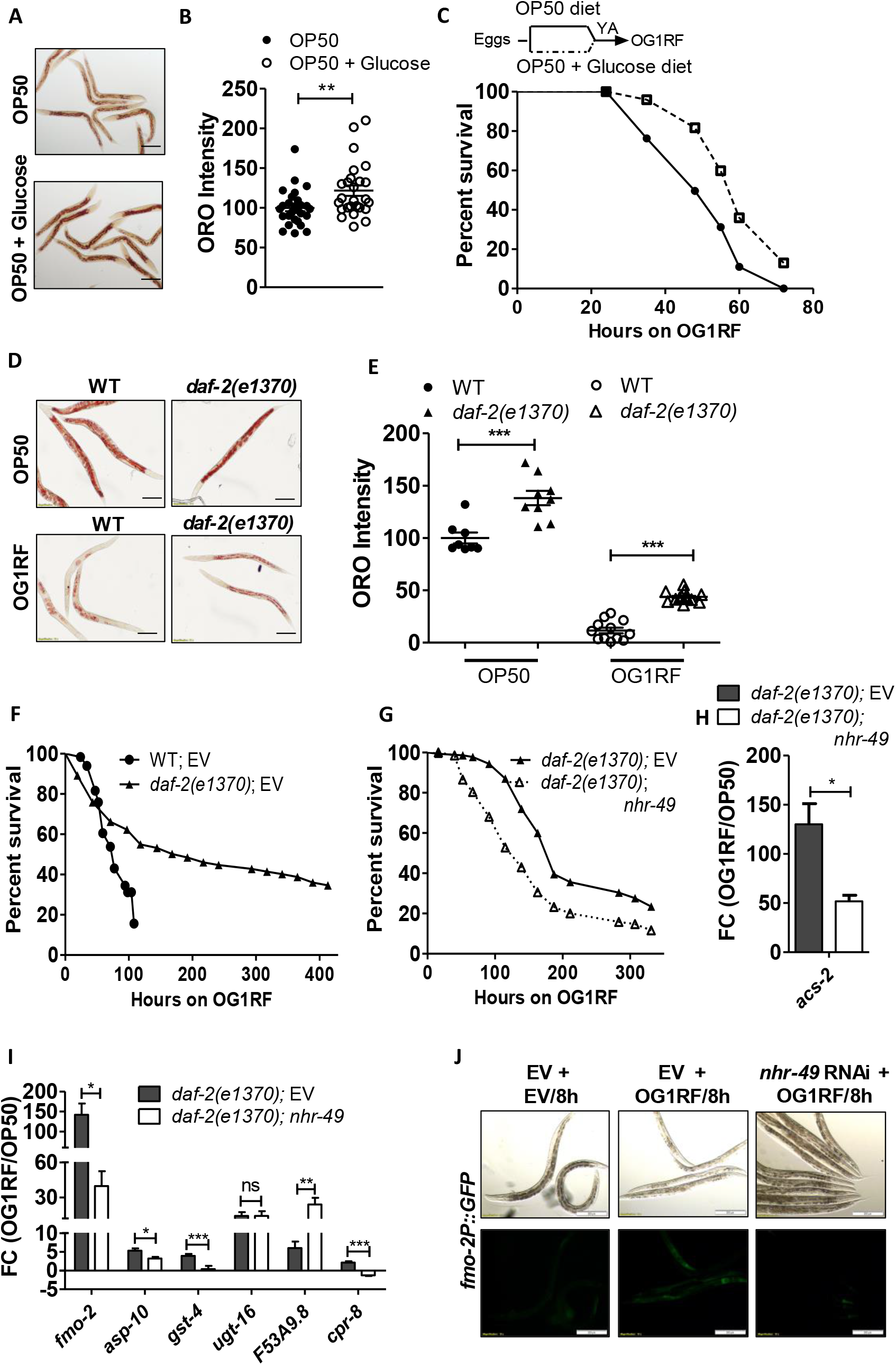
Lipids boost *C. elegans* survival during *E. faecalis* infection. (A-B) ORO staining and quantification of adult *C. elegans* fed on OP50 + 10 mM glucose diet as compared to normal OP50 diet. (C) Kaplan Maier survival curve on OG1RF for animals from glucose supplemented diet and normal diet (P <0.0001). (D and E) ORO staining and quantification of stain WT and *daf-2(e1370)* animals fed OG1RF for 8h. (F) Kaplan Maier survival curve for WT and *daf-2(e1370)* animals exposed to OG1RF (P <0.0001). (G) Kaplan Maier survival curve for *daf-2(e1370); EV* and *daf-2(e1370); nhr-49* RNAi animals exposed to OG1RF (P <0.0001).(H) Quantitative real time PCR analysis of OG1RF induced expression of immune effectors in *daf-2(e1370)* animals with *nhr-49* RNAi. (J) OG1RF induced expression of *fmo-2P::GFP* in WT and *nhr-49* RNAi animals.

*C. elegans* mutants with higher fat content, such as insulin receptor *daf-2*, is resistant to a number of biotic and abiotic stresses (Lee, Murphy and Kenyon, 2009). Mutation in *daf-2* leads to resistance to many microbial pathogens including OG1RF (Garsin et al., 2003; Singh and Aballay, 2006) and is also known to have higher lipid content (Ashrafi et al., 2003). We found that *daf-2(e1370)* animals had 40% more ORO stain than WT animals (Figure 6D). *daf-2(e1370)* animals also showed LD depletion upon OG1RF feeding indicating that starvation response is induced by *E. faecalis* in *C. elegans* irrespective of the genetic background. However, OG1RF fed *daf-2(e1370)* animals had more LD stain than OG1RF fed WT animals indicating that utilization of LDs may be need based (Figure 6D and 6E). As reported earlier, we found that *daf-2(e1370)* animals were highly resistant to OG1RF (Figure 6F).

To check if utilization of LDs contributed to enhanced resistance in high fat animals, we studied the effect of *nhr-49* RNAi on enhanced resistance of *daf-2(e1370)* animals. We found that *daf-2(e1370)* animals with *nhr-49* RNAi were more susceptible to death by OG1RF than animals with control RNAi (Figure 6G). Also, *nhr-49* knockdown dampened the induction of *acs-2* in *daf-2(e1370)* animals fed OG1RF diet (Figure 6H). Next, we tested whether NHR-49 activity regulates immune effector expression in *daf-2(e1370)* animals. Indeed, *nhr-49* RNAi dampened the expression of *fmo-*2, *asp-10, cpr-8* and *gst-4* transcripts in *daf-2(e1370)* animals compared to vector control (Figure 6I). There was an increase in F53A9.8 expression but no change in expression of *ugt-16* transcript upon knockdown of *nhr-49* in *daf-2(e1370)* animals. *daf-2(e1370)* animals had higher basal level of *fmo-2* transcript, but wild type levels of *asp-10, cpr-8, F53A9.8, ugt-16* and *gst-4* compared to WT animals (Figure S9). Using *fmo-2P::GFP* transgenic animals, we showed that *nhr-49* was necessary for OG1RF mediated induction of *fmo-2* expression (Figure 6J). Thus, our experiments show that NHR-49 contributes to enhanced resistance of *daf-2(e1370).* NHR-49 regulates both metabolic response and immune effector dependent protective response *in daf-2* animals fed on *E. faecalis.* Taken together, our data suggests that increase in utilizable neutral lipid stores of *C. elegans* enhances its survival on *E. faecalis* in an NHR-49 dependent manner.

### Metabolic response to *E. faecalis* diet is conserved across rhabditids

Gram positive cocci are common in nematode habitats and it is conceivable that starvation response to this diet may have evolved in other nematodes. We tested the effect of OG1RF diet on two additional members of family rhabditidae-*Caenorhabditis briggsae* and *Pristionchus pacificus* (Kiontke, Fitch and York, 2005). OG1RF feeding caused 50% depletion of lipid stores in both the nematodes (Figure 7A, B, C and D).

**Figure 7.**
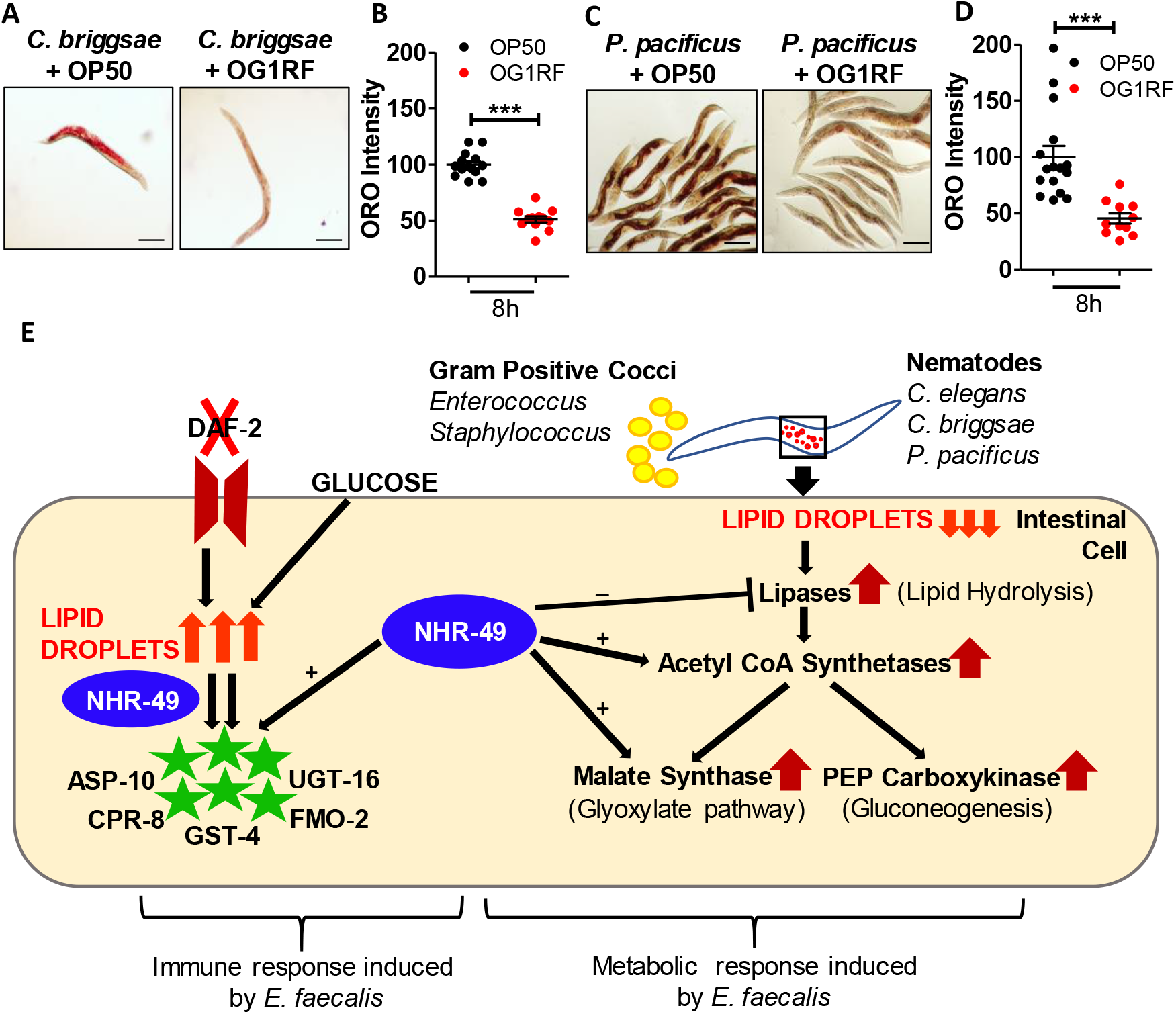
Metabolic response to *E. faecalis* diet is conserved across rhabtidae. (A-B) ORO staining and quantification of lipid droplets in adult *C. briggsae* AFI6 fed on OG1RF for 8h. (C-D) ORO staining and quantification of lipid droplets in adult *Pristionchus pacificus* PS312 fed on OG1RF for 8h. (E) Model: *C. elegans* fed on *Enterococcus faecalis* undergoes lipid depletion within 8 hours, along with a concomitant increase in expression of genes involved in lipid hydrolysis, beta oxidation, glyoxylate shunt and gluconeogenesis. NHR-49 positively regulates different steps in the lipid breakdown pathway as well as immune effectors production, and is crucial for survival on OG1RF. Increased lipid droplets via glucose and/or in *daf-2* mutant provides resistance towards infection.

In addition, OG1RF diet also caused larval arrest when fed to L1 larvae of *P. pacificus* and *C. briggsae* (data not shown). Taken together, our data provides strong evidence that OG1RF feeding induces starvation like response in three phylogenetically diverse nematodes. We propose a model (Figure 7E) in which a diet of *E. faecalis* coccoid cells causes a metabolic response in *C. elegans* and other rhabditids. The metabolic response is partly orchestrated by nuclear hormone receptor NHR-49 which leads to upregulation of some of the enzymes involved in sequential breakdown of neutral lipids, and acetyl CoA utilization via glyoxylate shunt and gluconeogenesis. NHR-49 also regulates production of many immune effectors namely, FMO-2, CPR-8, ASP-10, UGT-16 and GST-4. The NHR-49 regulated immuno-metabolic axis is also functional in high fat mutant and contributes to its enhanced resistance.

## DISCUSSION

### Gram positive cocci diet induces a metabolic response in *C. elegans* during early hours of infection

In this study, we show that two Gram positive cocci induce a ‘starvation like’ metabolic response in *C. elegans* which is not observed during infection with Gram negative bacteria. *E. faecalis*, a Gram positive coccoid bacterium, induces rewiring of the metabolic program in *C. elegans* which impinges on neutral lipid breakdown. This is accompanied by activation of glyoxylate shunt and of gluconeogenesis pathway, suggesting that stored lipids are likely utilized for sustenance in *E. faecalis* induced starvation. At least 25 genes regulating lipid breakdown are induced between 3 to 60 fold during *E. faecalis* exposure. We found evidence for metabolic rewiring in a recent study of *C. elegans* fed on *E. faecalis* strain MMH594 (Yuen and Ausubel, 2018) and also on *Staphylococcus* aureus. At least 7 transcripts encoding lipases, carnitine palmitoyl transferases, enoyl CoA hydratases enzymes directly involved in lipid breakdown were upregulated in *S. aureus* fed animals (Irazoqui et al., 2010). *P. aeruginosa* fed animals had more than 4 fold upregulation of transcript for *ech-9* and *cpt-4*, however *acs-2* transcript was only upregulated 3 fold (Troemel et al., 2006) compared to ∼40 fold on *E. faecalis*. Although we can not rule out utilization of lipid resources in other infection at a later time point, our study suggests that lipid utilization is a conserved response of nematodes to Gram positive coccoid bacteria.

It is conceivable that starvation response to *E. faecalis* (and to *S. aureus*) arises from inability of *C. elegans* to digest the cell wall of these bacterial cells. *C. elegans* genome encodes 13 lysozymes to digest peptidoglycan cell wall of bacteria. Of these, transcripts for 3 lysozymes-*ilys-2, lys-4* and *lys-7* are upregulated 4-20 fold in animals fed *E. faecalis* (Table S1). Lysozyme is a muramidase which cleaves glycosidic linkage in the peptidoglycan layer of bacterial cell wall and resistance to lysozyme can come from modification of sugars in the peptidoglycan layer (Davis and Weiser, 2011).*E. faecalis* strains, as well as *S. aureus* are known to be resistant to lysozyme (Hébert et al., 2007a; Herbert et al., 2007b) and this may be one reason for the inability of *C. elegans* to digest them, hence further contributing to starvation response.

### NHR-49 nuclear hormone receptor regulates both metabolic and immune response to *E. faecalis* infection

Lipid homeostasis comprising of lipid breakdown and synthesis is a highly regulated process. A number of molecular players as SREBP (SBP-1), PPARα (NHR-49), TOR, AMPK and hexosamine pathways are involved. Nuclear hormone receptor NHR-49, is a key regulator of the metabolic response induced in *C. elegans* during starvation (Van Gilst *et al.*, 2005; Van Gilst, Hadjivassiliou and Yamamoto, 2005). Other NHRs regulate fatty acid synthesis positively (NHR-80) or negatively (NHR-64) (Liang et al., 2010; Toselli-mollereau et al., 2011). Also, many NHRs which show upregulation during starvation followed by down regulation upon refeeding, suggesting that NHRs can modulate starvation response (Hyun et al., 2016). Phenotypically, *nhr-49* deletion mutants have increased fat stores, and it can regulate as many as 13 metabolic response genes involved in beta-oxidation, fatty acid desaturation, fatty acid binding/transport and glyoxylate pathways during starvation (Van Gilst, Hadjivassiliou and Yamamoto, 2005). In our study, we report a novel role of NHR-49 in metabolic regulation during *E. faecalis* diet. Of the 38 lipid metabolism genes dysregulated upon feeding on *E. faecalis*, at least seven are regulated by NHR-49. NHR-49 is a positive regulator of enzymes involved in TAG breakdown (*lipl-3*), fatty acid beta oxidation (*acs-2)* as well as glyoxylate shunt (*icl-1).* It also appears to be a negative regulator of *lipl-1, lipl-2, fil-1* and *ech-9*. This suggests that there might be an alternate mechanism of gene regulation for *lipl-1, lipl-2, fil-1* and *ech-9* which may be suppressed by *nhr-49*. Similar metabolic axis is known for PPARα in higher organisms such as mouse (Leone, Weinheimer and Kelly, 1999). We believe that alternate mechanisms of lipid hydrolysis also exist in *C. elegans* and should be examined in future studies.

Effect of NHR-49 knockdown on susceptibility to *E. faecalis* infection has been reported earlier although the mechanism was not studied (Sim and Hibberd, 2016). To our surprise, we find that NHR-49 regulated two distinct set of effectors-(a) lipid metabolic enzymes and (b) immune effectors such as aspartyl protease (*asp-10*), cysteine protease (*cpr-8*) and UDP-glycosyl transferase (*ugt-16*), glutathione-S-transferase *gst-4* and flavin mono-oxygenase (*fmo-2)*. It has been reported that NHR-49 activity positively regulates expression of detoxification genes such as glutathione-S-transferase (*gst-4*). While we were writing the manuscript another study indicated that flavin mono-oxygenase (*fmo-2*) upregulation during oxidative stress was dependent on NHR-49 (Qi et al., 2017; Goh et al., 2018; Hu et al., 2018). Our study shows that NHR-49 regulates detoxification enzymes in the context of infection as well. It was interesting to note that enhanced resistance phenotype of insulin receptor mutant *daf-2* towards *E. faecalis* was partially dependent on NHR-49, through its effect on the expression of immune effectors. We propose that lipid droplets are used by *C. elegans* to derive energy to generate immune effectors. Two instances of increased neutral lipids, *daf-2* mutation and glucose supplementation, increased *C. elegans* survival on *E. faecalis*. This supports our hypothesis that lipid droplets have a protective role to play during infection condition. To the best of our knowledge, we provide NHR-49 as the first example of a metabolic regulator engaging in immune effector production.

There are a number of examples of LD dys-homeostasis during pathogenesis outside nematodes. Bacterial pathogens such as *Chlamydia trachomatis* and *Mycobacterium tuberculosis* require the presence of host LDs for their survival (Kumar, Cocchiaro and Valdivia, 2006; Daniel et al., 2011). Under hypoxia, Mycobacterium infection induces the accumulation of LDs in immune cells to form foamy macrophage. This bacterium can scavange fatty acids from the host LDs and accumulates triacylglycerol in its cytosol which promotes dormancy (Daniel et al., 2011). In Hepatitis C Virus infection, host LDs can also be used as platforms for viral assembly (Miyanari et al., 2007; Boulant, Targett-Adams and Mclauchlan, 2007) while LDs bound to histones are proposed to be a conserved mechanism of immune response (Anand et al., 2012). In all these studies suggest that energy stored in lipid droplets can be utilized by the pathogens or by the host, and can cause a tug of war leading to metabolic rewiring.

## CONCLUSION

Our data suggests that energy derived from the lipid droplets is used for boosting resistance mechanisms and hence survival. Recent reports from animal models of infection show that host can withhold macronutrients as well as important transition metal ions such as iron, zinc, copper and manganese from the pathogen to reduce its replication. This process is known as Nutritional Immunity (Hood and Skaar, 2012). Our study presents a new facet of nutritional immunity wherein, the host utilizes its own nutritional reservoirs to boost immune response and survival. This study also highlights the possible therapeutic role of lipids and fats during infection. A detailed study of how LD levels are regulated during microbial infection of different kinds will be a goal in the near future.

## STAR METHODS

### Bacterial strains and Growth media

All bacterial strains used in this study are listed in Table S1. *E. coli* was grown in LB at 37°C for 8 hours and seeded on NGM plates with streptomycin. For making pathogen lawns, *E. faecalis* strains were grown in BHI with appropriate antibiotics for 5 hours at 37°C and 50μl was spread on BHI agar plates with respective antibiotics and kept at 37°C overnight. Gentamycin (50μg/ml) was used for OG1RF, erythromycin (12.5μg/ml) and tetracycline (4μg/ml) was used for OG1RF::GFP, Rifampicin (100μg/ml) was used for TX5266 *ΔfsrB* OG1RF and erythromycin (10μg/ml) for TX5266.01 complemented strain of OG1RF. For experiments, OG1RF was seeded on NGM plates. *S. aureus* was grown in BHI broth supplemented with 5μg/ml nalidixic acid followed by plating on BHI agar plate. *P. aeruginosa* was grown in LB for 8-10 hours and spread on SK agar plate (Singh and Aballay, 2006). *S. typhimurium* was grown for 4-5 hours in LB and spread on NGM plates.

### *C. elegans* strains

N2 (Bristol) was used as *C. elegans* wildtype strain and animals were maintained with *E. coli* OP50 on Nematode Growth Medium (NGM), as previously described (Brenner, S., 1974). All strains used in this study are listed in Table 1. All animals have been maintained at 20°C except for temperature sensitive mutant *daf-2(e1370)*, which has been maintained at 15°C. All experiments were conducted at 25°C.

### RNA interference

The *C. elegans* RNAi library was obtained from Source BioScience. Individual HT115(DE3) bacteria clones expressing dsRNA against genes of interest were grown at 37°C in LB with carbenicillin (25μg/ml), then seeded onto NGM-carbenicillin plates supplemented with 2.5mM IPTG. Adult animals were allowed to lay eggs on RNAi plates and allowed to develop till young adult. Animals were transferred to *E. faecalis* lawn prior to lipid staining, RNA extraction or survival assay.

### Body length measurements

Synchronized L1 larvae (Figure 2) or adults (Figure 3) were exposed to OP50 and OG1RF lawns. After every 24 hours 15-20 animals were mounted on 2% agarose and imaged. Using AxioCam I Cm1, Zeiss. Body length of animals was measured using Image J. Each experiment was repeated at least three times.

### Vulva and Gonad Development

For vulva development studies, PS3525, *egl-17::cfp + unc-119(+)* was used to trace vulval precursor cell lineage during development (Figure S2). Synchronized L1s of PS3525 were exposed to OP50 and OG1RF and imaged at 20h and 84h post exposure using LSM880 airyscan microscope. The number of cells showing *egl-17::cfp* fluorescence were counted. 10-20 animals were used for each time point for each diet.

For tracing gonad development, synchronized wild type L1 larvae were grown on OP50 and OG1RF lawn and 10-15 animals were imaged for each condition. Gonad proliferation is described (Fig S3). Number of cells in the gonadal primordium were visualized and counted using differential interference contrast (DIC) microscopy at 12h, 36h and 60h of exposure of animals to OP50 and OG1RF diet.

### *C. elegans* survival assay

*E. faecalis* lawns were made by spreading 50μl of bacterial culture on 60mm BHI agar plates and incubated at 37°C for 12-16 hours. In some cases, *E. faecalis* lawns were made on NGM agar plates as described. 100-120 synchronized young adult animals were exposed to pathogen at 25°C, scored for survival at the times indicated in the survival assay graphs. Animals were considered dead when they failed to respond to touch. Each survival assays was performed three times or more.

### Quantification of Intestinal Bacterial Loads and Lumen Distension

Quantification of intestinal bacterial loads was done as mentioned (Singh and Aballay, 2012). Briefly, synchronized adult animals exposed to OG1RF GFP, were transferred to a fresh OP50 plate and allowed to crawl for few minutes to eliminate OG1RF stuck to cuticle. Groups of 10 animals were washed three times and then crushed in 50ul PBS containing 0.1% Triton X-100 to release proliferating bacteria in the intestine. Serial dilutions of the lysates were plated on BHI agar plates with erythromycin and tetracycline antibiotics to select for OG1RF-GFP cells. Single colonies were counted next day and represented as colony forming units (cfu) per animal. Seven independent sets of 10 animals each were used for cfu analysis.

For measuring intestinal distention adult animals fed on OP50 and OG1RF, were mounted on fresh agarose pad slides with a drop of 1ul sodium azide (1mM) in 10ul M9 buffer and imaged using AiryScan microscope at 63X.

### Carbohydrate and Protein Estimation

Carbohydrate content in *C. elegans* was estimated by the Anthrone assay as described (Morris, 1948). Briefly, 800-1000 animals, exposed to OP50 and OG1RF for 8 hours, were collected in in tubes using M9 buffer and sonicated (55% amplitude, pulse on − 1min, pulse off − 30seconds) for 45 minutes using Branson digital sonifier. To 100μl of the sonicated lysate 500μl of 0.14% Anthrone reagent was added and the mixture was incubated at 80°C for 20 minutes. The absorbance was read at 620nm using Infinte M200pro TECAN machine. The amount of carbohydrate content per mL of the sample and per animals was calculated based on a glucose standard curve prepared using anthrone reagent. Protein estimation was done using Bradford Assay (Bradford, 1976). To 20ul sonicated lysate, 40ul of Bradford reagent and 240ul of water was added. The reaction was incubated in dark for 30 minutes and absorbance was measured at 595nm. Protein content of animals was estimated by the Bradford assay against a standard curve of bovine serum albumin. Carbohydrate and protein estimation tests were done three independent times.

### Oil Red O staining and Quantification

Animals were stained with Oil Red O as described in (Rourke, Soukas and Carr, 2010). Briefly, synchronized population of adult *C. elegans*, on different diets, were fixed by incubating the animals with equal volumes of PBS buffer and 2X MRWB (160mM KCl, 40mM NaCl, 14mM Na^2^EGTA, 1mM spermidine-HCl, 0.4mM spermine, 30mM Na-PIPES (pH 7.4), 0.2% beta mercaptoethanol and 20mg/ml PFA) for 1 hour at room temperature. Animals were washed three times with PBS and then incubated with 60% isopropanol for 15 minutes at room temperature. Prior to adding the stain to the animals, 60% ORO was freshly prepared in water and kept for rotation for 1 hour and then filtered. Animals were incubated with the stain for 16-18 hours, following which they were washing with PBS + 0.1% triton × 100. Animals were mounted on agarose pads for imaging under Olympus IX81 bright field microscope. Images were quantified using ImageJ software. All the experiments were performed at least 3 times. ORO stain was quantified in 15-20 animals in each experimental set/strain/treatment.

### BODIPY Staining and Quantification

Animals were stained as described (Klapper, 2011). Synchronized population of adult animals were fixed in 4% paraformaldehyde followed by three cycles of freeze thaw. Fixed animals were stained in 1µg/ml BODIPY ^493/503^ in M9 buffer, for 1 hour at room temperature. The stained animals were imaged using Olympus IX81 fluorescence microscope. Image analysis and quantification was done using ImageJ.

### Fluorescence Microscopy

Adult *acs-2P::GFP* and adult *fmo-2P::GFP* animals were allowed to feed on OP50 and OG1RF for 8 hours. 10-12 animals were mounted on fresh agarose-pad slides and imaged using fluorescence microscope Olympus IX81. Each experiment was done four times or more.

### Real time PCR

800-1000 synchronized adult animals were exposed to OP50 and OG1RF for 8 hours. Animals were collected using M9 buffer and frozen in QIAzol lysis reagent at −80°C. Total RNA was extracted using RNeasy Plus Universal Mini Kit (QIAGEN), and reverse transcribed using iScript cDNA synthesis kit (BIORAD). cDNA was subjected to qRT PCR analysis using SYBR Green detection (iTaq Universal supermix, BIORAD) on QuantStudio3 (Applied Biosystems) machine. Primers for qRT PCR were designed using Primer 3 online software. All Ct values were normalized using control gene (*actin-1*) *act-1*. Comparative ΔCt method was used to determine fold change in gene expression (Livak and Schmittgen, 2001). All experiments were repeated at least three times and fold change data was presented as mean ±sem. Primer sequences are available upon request.

### RNA-sequencing and Data Analysis

RNA was isolated using Qiagen Universal RNA isolation kit from approximately 1000 L4 animals exposed to OP50 and OG1RF for 8 hours. Three biological replicates were prepared for each. cDNA library was prepared using NEBNext Ultra II Directional RNA Library Prep Kit for Illumina. The libraries were subjected to 50-base pair single-end sequencing using Illumina sequencer (services from Genotypic Inc). The data was analysed using the following pipeline (Pertea *et al.*, 2016). HISAT was used to map the RNA-seq reads onto the latest *C. elegans* reference genome (WS266) from WORMBASE resulting in a Sequence Alignment Map (SAM). SAM format was converted to its binary form BAM using SAMTOOLS. BAM files were then used as input for assembling the transcripts using STRINGTIE. Stringtie was used to assemble and quantify the levels of expressed genes to produce Fragments Per Kilobase of exon per Million fragments mapped (FPKM). The assembly was merged into a singular Gene Transfer Format (GTF) to facilitate comparison with the reference annotation in the same format. BALLGOWN was used to calculate differential expression of genes using FPKM data and produces table with fold change and P-value and q-value. Genes were shortlisted with a cutoff of 1.5 fold change and P-values less than 0.05.

### Statistical analysis

Survival curves for *C. elegans* adults were plotted using the Graphpad PRIZM. Prism uses the Kaplan-Meier method to calculate survival fractions and the log rank test to compare survival curves. Survival curves were considered different from the appropriate control when p values were < 0.05. A two-sample *t* test for independent samples was used to analyse oil red O, BODIPY, body measurements, protein and carbohydrate analysis, cfu counts, fluorescence intensity experiments and qRT-PCR results. p values < 0.01 are considered significant. All experiments were repeated at least three times unless otherwise indicated.

## ACKNOLEDGMENTS

We thank Dr Barbara E Murray, University of Texas Health Science Centre at Houston and Prof Manuel Espinosa, Spanish National Research Council for *Enterococcus faecalis* strains. Some *C. elegans* strains were provided the CGC which is funded by the NIH Office of Infrastructure Programs (P40 OD01440). MD is supported by scholarship from University Grants Commission of the Government of India. This work was supported by the Wellcome Trust/DBT India Alliance Intermediate Fellowship (Grant no. IA/I/13/1/500919) awarded to Varsha Singh.

## AUTHOR CONTRIBUTIONS

VS and MD conceptualized the project. MD, MS, NB and SJ performed the experiments. VS, MD, MS and NB analysed the data and interpreted the results. AG performed RNA sequencing analysis. VS, MD, NB and AG wrote the manuscript.

## REFERENCES

Anand, P., Cermelli, S., Li, Z., Kassan, A., Bosch, M., Sigua, R., Huang, L., Ouellette, A.J., Pol, A., Welte, M.A. and Gross, S.P., 2012. A novel role for lipid droplets in the organismal antibacterial response. Elife, 1, p.e00003.

Ashrafi, K., Chang, F.Y., Watts, J.L., Fraser, A.G., Kamath, R.S., Ahringer, J. and Ruvkun, G., 2003. Genome-wide RNAi analysis of Caenorhabditis elegans fat regulatory genes. Nature, 421(6920), pp.268–272

Baugh, L.R., 2013. To grow or not to grow: Nutritional control of development during Caenorhabditis elegans L1 Arrest. Genetics, 194(3), pp. 539–555.

Boulant, S., Targett-Adams, P. and McLauchlan, J., 2007. Disrupting the association of hepatitis C virus core protein with lipid droplets correlates with a loss in production of infectious virus. Journal of General Virology, 88(8), pp.2204–2213.

Bradford, M.M., 1976. A rapid and sensitive method for the quantitation of microgram quantities of protein utilizing the principle of protein-dye binding. Analytical biochemistry, 72(1-2), pp.248–254.

Brenner, S., 1974. The genetics of Caenorhabditis elegans. Genetics, 77(1), pp.71–94.

Burdine, R.D., Branda, C.S. and Stern, M.J., 1998. EGL-17(FGF) expression coordinates the attraction of the migrating sex myoblasts with vulval induction in C. elegans. Development 125(6), pp. 1083–93.

Cassada, R.C. and Russell, R.L., 1975. The dauerlarva, a post-embryonic developmental variant of the nematode Caenorhabditis elegans. Developmental Biology, 46(2), pp. 326–342.

Daniel, J., Maamar H., Deb, C., Sirakova, T.D. and Kolattukudy P.E., 2011. Mycobacterium tuberculosis uses host triacylglycerol to accumulate lipid droplets and acquires a dormancy-like phenotype in lipid-loaded macrophages. PLoS Pathogens, 7(6), e1002093

Davis, K.M. and Weiser, J.N., 2011. Modifications to the peptidoglycan backbone help bacteria to establish infection. Infection and Immunity, 79(2), pp. 562–570.

Engelmann, I., Griffon A., Tichit, L., Montanana-Sanchis, F., Wang G., Reinke, V., Waterstone R.H., Hillier L.W. and Ewbank, J.J., 2011. A comprehensive analysis of gene expression changes provoked by bacterial and fungal infection in C. elegans. PLoS ONE, 6(5), e19055

Garsin, D.A., Sifri, C.D., Mylonakis, E., Qin, X., Singh, K.V., Murray, B.E., Calderwood, S.B. and Ausubel, F.M., 2001. A simple model host for identifying Gram-positive virulence factors. Proceedings of the National Academy of Sciences, 98(19), pp.10892–10897.

Garsin, D.A., Villanueva, J.M., Begun, J., Kim, D.H., Sifri, C.D., Calderwood, S.B., Ruvkun, G. and Ausubel, F.M., 2003. Long-lived C. elegans daf-2 mutants are resistant to bacterial pathogens. Science, 300(5627), p.1921

Goh, G.Y.S., Winter, J.J., Bhanshali, F., Doering, K.R.S., Lai, R., Lee, K., Veal, E.A. and Taubert, S., 2018. NHR-49/HNF 4 integrates regulation of fatty acid metabolism with a protective transcriptional response to oxidative stress and fasting. Aging cell, 17(3), p.e12743

Golden, J.W. and Riddle, D.L., 1982. A pheromone influences larval development in the nematode Caenorhabditis elegans. Science, 218(4572), pp.578–580.

Han, S., Schroeder, E.A., Silva-García, C.G., Hebestreit, K., Mair, W.B. and Brunet, A., 2017. Mono-unsaturated fatty acids link H3K4me3 modifiers to C. elegans lifespan. Nature, 544(7649), pp.185–190.

Goudeau, J. Bellemin, S., Toselli-mollereau, E., Shamalnasab, M., Chen, Y. and Aguilaniu H., 2011. Fatty Acid Desaturation Links Germ Cell Loss to Longevity Through NHR-80 / HNF4 in C. elegans’. PLoS Biology, 9(3), p.e1000599.

Hart, B.L., 1988. Biological basis of the behavior of sick animals. Neuroscience & Biobehavioral Reviews, 12(2), pp.123–137.

Hébert, L., Courtin, P., Torelli, R., Sanguinetti, M., Chapot-Chartier, M.P., Auffray, Y. and Benachour, A., 2007a. Enterococcus faecalis constitutes an unusual bacterial model in lysozyme resistance. Infection and immunity, 75(11), pp.5390–5398.

Herbert, S., Bera, A., Nerz, C., Kraus, D., Peschel, A., Goerke, C., Meehl, M., Cheung, A. and Götz, F., 2007b. Molecular basis of resistance to muramidase and cationic antimicrobial peptide activity of lysozyme in staphylococci. PLoS pathogens, 3(7), p.e102.

Hood, M.I. and Skaar, E.P., (2012) ‘Nutritional immunity?: transition meatls at the pathogen-host interface’, Nature Reviews. Microbiology, 10(8), pp. 525–537

Hu, Q., D’Amora, D.R., MacNeil, L.T., Walhout, A.J. and Kubiseski, T.J., 2018. The Caenorhabditis elegans Oxidative Stress Response Requires the NHR-49 Transcription factor. G3(Bethesda), 8(12), pp.3857–3863

Hyun, M., Davis, K., Lee, I., Kim, J., Dumur, C. and You, Y.J., 2016. Fat metabolism regulates satiety behavior in C. elegans. Scientific reports, 6, p.24841.

Irazoqui, J.E., Troemel, E.R., Feinbaum, R.L., Luhachack, L.G., Cezairliyan, B.O. and Ausubel, F.M., 2010. Distinct pathogenesis and host responses during infection of C. elegans by P. aeruginosa and S. aureus. PLoS pathogens, 6(7), p.e1000982.

Kimble, J. and Hirsh, D.,1979. The postembryonic cell lineages of the hermaphrodite and male gonads in Caenorhabditis elegans. Developmental Biology, 70(2), pp. 396–417.

Kiontke, K. and Fitch, D.H., 2005. The phylogenetic relationships of Caenorhabditis and other rhabditids. WormBook, 11, pp.1–11.

Klapper, M., Ehmke, M., Palgunow, D., Böhme, M., Matthäus, C., Bergner, G., Dietzek, B., Popp, J. and Döring, F., 2011. Fluorescence-based fixative and vital staining of lipid droplets in Caenorhabditis elegans reveal fat stores using microscopy and flow cytometry approaches. Journal of lipid research, 52(6), pp.1281–1293.

Kumar, Y., Cocchiaro, J. and Valdivia, R.H., 2006. The obligate intracellular pathogen Chlamydia trachomatis targets host lipid droplets. Current biology, 16(16), pp.1646–1651.

Larance, M., Pourkarimi, E., Wang, B., Murillo, A.B., Kent, R., Lamond, A.I. and Gartner, A., 2015. Global proteomics analysis of the response to starvation in C. elegans. Molecular & Cellular Proteomics, 14(7), pp.1989–2001.

Lee, K.A., 2006. Linking immune defenses and life history at the levels of the individual and the species. Integrative and comparative biology, 46(6), pp. 1000–1015.

Lee, S.J., Murphy, C.T. and Kenyon, C., 2009. Glucose Shortened the Lifespan of Caenorhabditis elegans by Down-Regulating Aquaporin Gene Expression. Cell Metabolism. 10(5), pp. 379–391.

Liang, B., Ferguson, K., Kadyk, L. and Watts, J.L., 2010. The role of nuclear receptor NHR-64 in fat storage regulation in Caenorhabditis elegans. PloS one, 5(3), p.e9869.

Leone, T.C., Weinheimer C.J. and Kelly, D.P.,1999. A critical role for the peroxisome proliferator-activated receptor alpha (PPARalpha) in the cellular fasting response?: The PPARalpha-null mouse as a model of fatty acid oxidation disorders. PNAS. 96(June), pp. 7473–7478.

Liu, F., Xiao, Y., Ji, X.L., Zhang, K.Q. and Zou, C.G., 2017. The cAMP-PKA pathway-mediated fat mobilization is required for cold tolerance in C. elegans. Scientific reports, 7(1), p.638.

Livak, K.J. and Schmittgen, T.D., 2001. Analysis of relative gene expression data using real-time quantitative PCR and the 2− ΔΔCT method. Methods, 25(4), pp.402–408.

Lynn, D.A., Dalton, H.M., Sowa, J.N., Wang, M.C., Soukas, A.A. and Curran, S.P., 2015. Omega-3 and-6 fatty acids allocate somatic and germline lipids to ensure fitness during nutrient and oxidative stress in Caenorhabditis elegans. Proceedings of the National Academy of Sciences, 112(50), pp.15378–15383.

Miyanari, Y., Atsuzawa, K., Usuda, N., Watashi, K., Hishiki, T., Zayas, M., Bartenschlager, R., Wakita, T., Hijikata, M. and Shimotohno, K., 2007. The lipid droplet is an important organelle for hepatitis C virus production. Nature cell biology, 9(9), pp.1089–1097.

Morris, D.L., 1948. Quantitative determination of carbohydrates with Dreywood’s anthrone reagent. Science, 107, pp.254–255.

Nomura, T., Horikawa, M., Shimamura, S., Hashimoto, T. and Sakamoto, K., 2010. Fat accumulation in Caenorhabditis elegans is mediated by SREBP homolog SBP-1. Genes & nutrition, 5(1), pp.17–27.

O’Rourke, E.J., Soukas, A.A., Carr, C.E. and Ruvkun, G., 2009. C. elegans major fats are stored in vesicles distinct from lysosome-related organelles. Cell metabolism, 10(5), pp.430–435.

O’Rourke, E.J. and Ruvkun, G., 2013. MXL-3 and HLH-30 transcriptionally link lipolysis and autophagy to nutrient availability. Nature cell biology, 15(6), p.668.

Pertea, M. Kim, D., Pertea, G.M., Leek, J.T. and Salzberg, S.L., 2016. Transcript-level expression analysis of RNA-seq experiments with HISAT, StringTie and Ballgown. Nature Protocols, 11(9), pp. 1650–1667.

Qi, W., Gutierrez, G.E., Gao, X., Dixon, H., McDonough J.A., Marini, A.M. and Fisher, A.L., 2017. The ω-3 fatty acid α-linolenic acid extends *Caenorhabditis elegans* lifespan via NHR-49/PPARα and oxidation to oxylipins. Aging Cell, 16(5), pp. 1125–1135.

Qin, X., Singh, K.V., Weinstock, G.M. and Murray, B.E., 2000. Effects of Enterococcus faecalis fsr genes on production of gelatinase and a serine protease and virulence. Infection and Immunity, 68(5), pp. 2579–2586.

Ratnappan, R., Amrit, F.R., Chen, S.W., Gill, H., Holden, K., Ward, J., Yamamoto, K.R., Olsen, C.P. and Ghazi, A., 2014. Germline Signals Deploy NHR-49 to Modulate Fatty-Acid β-Oxidation and Desaturation in Somatic Tissues of C. elegans. PLoS Genetics, 10(12). e1004829

Rauw, W.M., 2012. Immune response from a resource allocation perspective. Frontiers in Genetics, pp. 1–14.

Sim, S. and Hibberd, M.L., 2016. Caenorhabditis elegans susceptibility to gut Enterococcus faecalis infection is associated with fat metabolism and epithelial junction integrity. BMC microbiology, 16:6

Singh, V. and Aballay, A., 2006. Heat-shock transcription factor (HSF)-1 pathway required for Caenorhabditis elegans immunity. Proceedings of the National Academy of Sciences, 103(35), pp.13092–13097.

Singh, V. and Aballay, A., 2012. Endoplasmic Reticulum Stress Pathway Required for Immune Homeostasis Is Neurally Controlled by Arrestin-1. J. Biol. Chem. 287(40), pp. 33191–33197.

Sulston, J.E. and Horvitz, H.R., 1977. Post-embryonic cell lineages of the nematode, Caenorhabditis elegans. Developmental Biology, 56(1), pp. 110–156.

Tang, R.J., Breger, J., Idnurm, A., Gerik, K.J., Lodge, J.K., Heitman, J., Calderwood, S.B. and Mylonaskis, E., 2005. Cryptococcus neoformans gene involved in mammalian pathogenesis identified by a Caenorhabditis elegans progeny-based approach. Infection and Immunity, 73(12), pp. 8219–8225.

Taubert, S., Van Gilst, M.R., Hansen, M. and Yamamoto, K.R., 2006. A mediator subunit, MDT-15, integrates regulation of fatty acid metabolism by NHR-49-dependent and -independent pathways in C. elegans. Genes and Development, 20(9), pp. 1137–1149.

Troemel, E.R., Chu, S.W., Reinke, V., Lee, S.S., Ausubel, F.M. and Kim, D.H., 2006. p38 MAPK regulates expression of immune response genes and contributes to longevity in C. elegans. PLoS genetics, 2(11), p.e183.

Vale, P.F., Fenton, A. and Brown, S.P., 2014. Limiting Damage during Infection : Lessons from Infection Tolerance for Novel Therapeutics. PloS Biology, 12(1), e1001769

Van Gilst, M. R., Hadjivassiliou, H. and Yamamoto, K. R., 2005. A Caenorhabditis elegans nutrient response system partially dependent on nuclear receptor NHR-49. Proceedings of the National Academy of Sciences of the United States of America, 102(38), pp. 13496–13501.

Van Gilst, M.R., Hadjivassiliou, H., Jolly, A. and Yamamoto, K.R., 2005. Nuclear hormone receptor NHR-49 controls fat consumption and fatty acid composition in C. elegans. PLoS biology, 3(2), p.e53.

Yuen, G.J. and Ausubel, F.M., 2018. Both live and dead Enterococci activate Caenorhabditis elegans host defense via immune and stress pathways. Virulence, 9(1), pp.683–699.

