## Supplementary Figures for "Nuclear hormone receptor NHR-49 shapes immuno-metabolic response of *Caenorhabditis elegans to Enterococcus faecalis* infection"

**FIGURE S1**

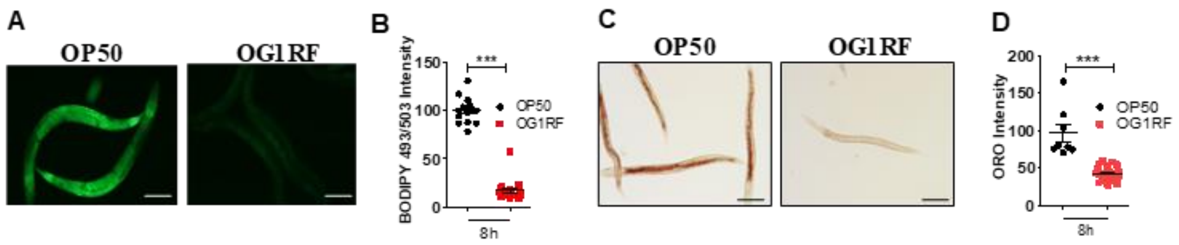


**Figure S1. *E. faecalis* induces lipid droplet depletion in *C. elegans*.** (A-B) BODIPY$^{493/503}$ staining and quantification of adult *C. elegans* fed on OG1RF for 8 hours. (C-D) ORO staining and quantification of young L4 larva fed on OG1RF for 8 hours. Scale bar, 100 μm.

FIGURE S2

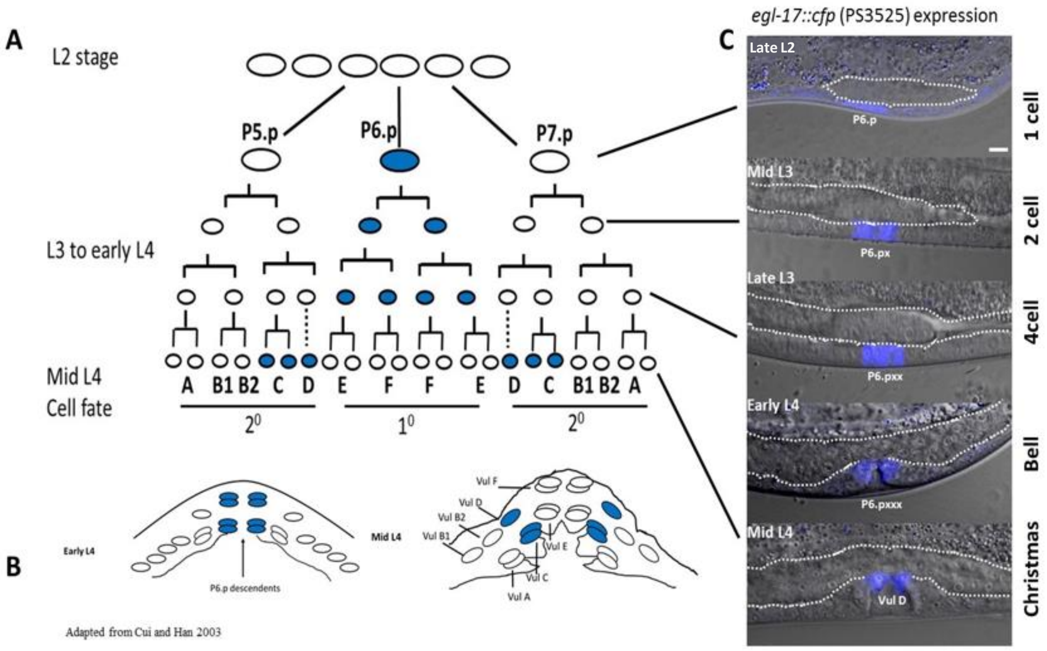


**Figure S2. Cell lineage analysis of vulva using *egl-17*::*cfp*.** (A) Cartoon showing vulval invariant cell lineages and morphogenesis in wild-type hermaphrodites (Burdine et al., 1998; Sulston and Horvitz, 1977). At larval stage, L3, vulval precursor cells P(5-7).p adopt primary (1°) or secondary (2°) cell fates and undergo invariant cell divisions to produce 22 cells organized into toroids, VulA, vulB1, vulB2, vulC, vulD, vulE, and vulF, shown by A, B1, B2, C, D, E, and F cell types (Sharma-Kishore et al., 1999). Blue colored oval cells indicate *egl-17::cfp* expression in specific lineages. (B) Cartoon showing early L4 vulva, when *egl-17::cfp* is exclusively seen in granddaughters of P6.p (1°), and mid-L4 stage, where the expression is shifted to 2° lineage cells, vulC and vulD. (C) Merged DIC and fluorescence images showing stage specific *egl-17::cfp* in vulval cells. P6.p (1 cell stage), P6.px (2 cell/midL3), P6.pxx (4 cell/mid to late L3), and P6.pxxx (8 cell/early L4 onwards), x denotes one round of mitotic division. White dashed shapes show gonad development. In all animals, anterior is to the left; scale bar, 5 µm.

FIGURE S3

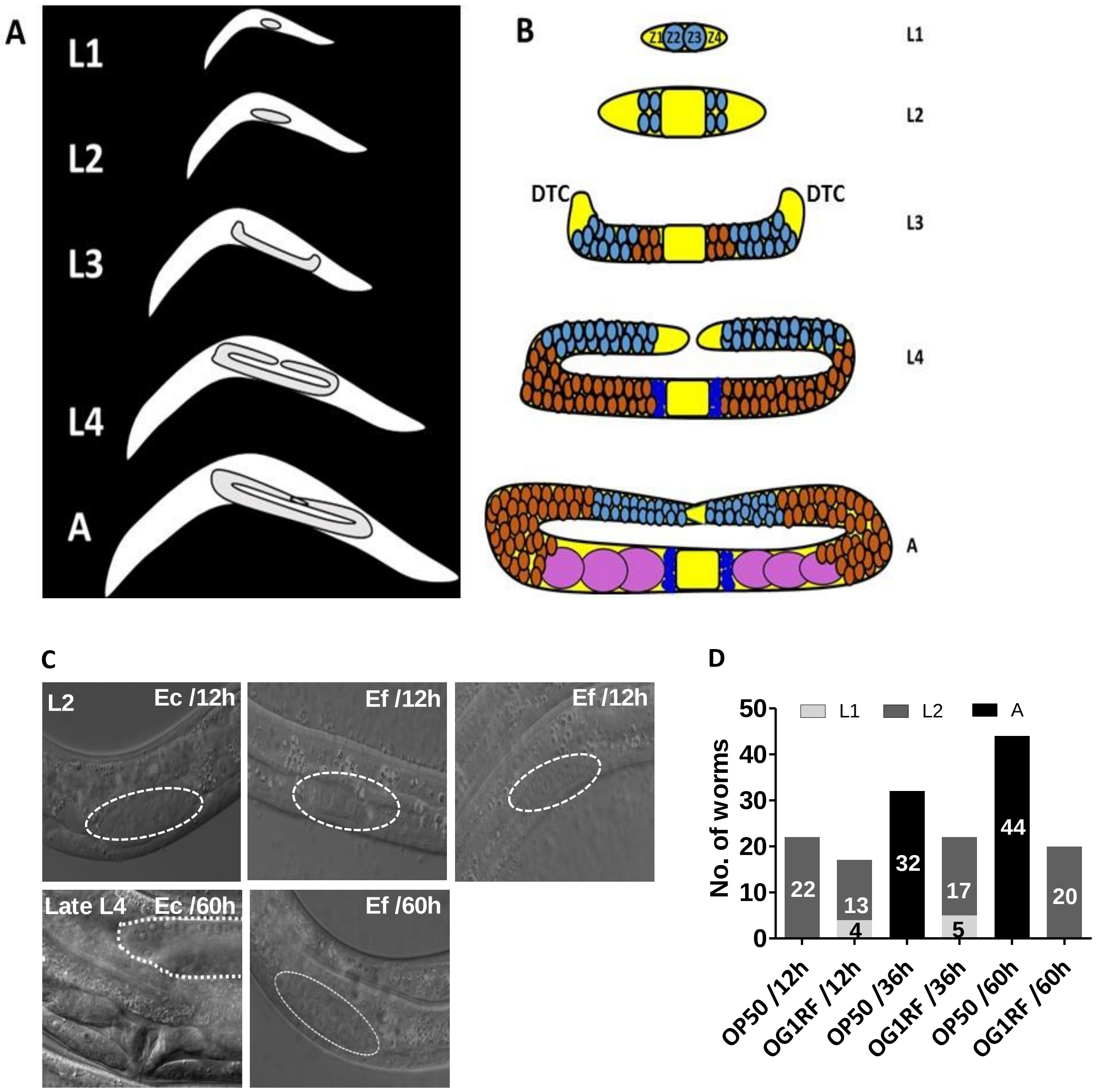


**Figure S3. Cell lineage analysis of gonad primordium in *E. faecalis* fed animals indicates L2 arrest.** (A) Schematic of shape changes in gonad primordium from L1 larva to adult *C. elegans* hermaphrodite. (B) Schematic of somatic gonad and germline proliferation from L1 to adulthood. At hatching, the gonad consists of 2 somatic precursor cells, Z1/Z4 and 2 germline founder cells, Z3/Z4. By late L2, Z1 and Z4 proliferate to produce 2 two distal tip cells (DTCs) and 10 proximal cells. At early L3, rapid germline proliferation occurs from Z2/Z3 followed by dorsally upward movement of gonad arms (mid L3) and further turning and extension towards vulva midline (mid L4). From late L4 to adulthood, both arms cross over the midline and overlap each other. DTCs and somatic gonad cells are yellow, the central oval cell depicts multiple somatic cells. Proliferating and meiotic S cells are in light blue, meiotic cells are brown and dark blue sperm and pink are oocytes. (C) Gonad development in OP50 fed animals is normal but fail to progress beyond 12 nuclei L2 stage in OG1RF fed animals. DIC images of gonad primordia in mid to late L2 and late L4 in OP50 fed animals. In OG1RF fed animals, either L1 or L2 gonad primordia were seen at 12h of feeding and only L2 primordia at 60h time point. Dashed white circles and shapes show gonad primordia. In all animals, anterior is to the left. Scale bar, 5 µm. (D) Quantification of stage specific gonad primordia in OP50 and OG1RF fed larvae. No of animals are indicated on the bars.

FIGURE S4

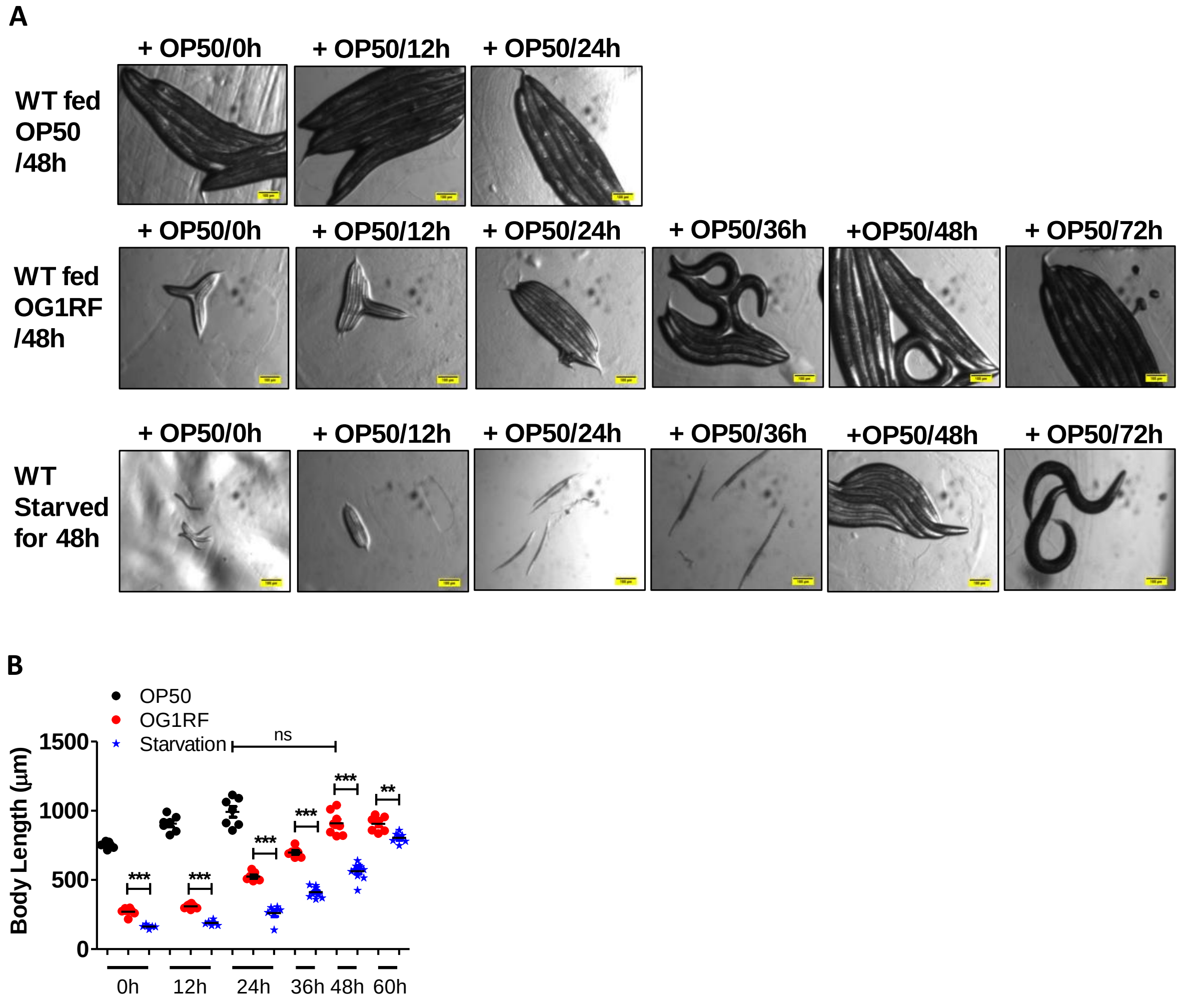


**Figure S4. *E. faecalis* induced L2 arrest in worms is reversible in nature.** (A) L1 larvae fed for 48 hours on OP50 or on OG1RF or starved for 48 hours were transferred to OP50 and imaged at different time points. (B) Mean ± SEM of body length of animals in panel A. Scale bar 100 µm.

FIGURE S5

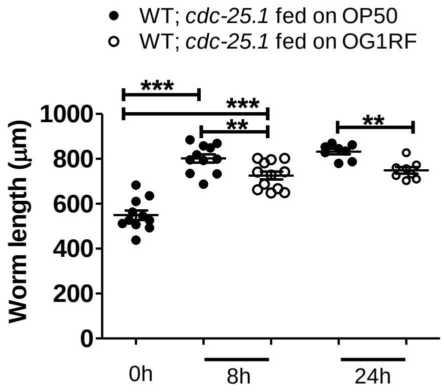


**Figure S5. *E. faecalis* diet leads to reduction in body size in adult without germline proliferation.** Scatter plot of body length in *cdc-25.1* RNAi animals fed OP50 or OG1RF.

FIGURE S6

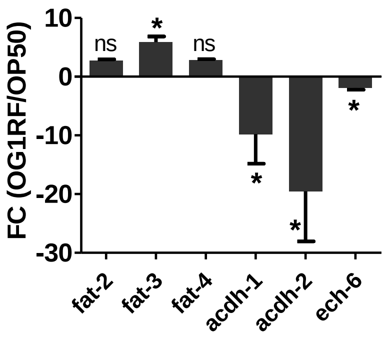


**Figure S6. *E. faecalis* diet causes dysregulation in the expression of genes involved in lipid homeostasis.** Quantitative real time PCR analysis (Mean ± SEM) of *fat-3*, *acdh-1, acdh-2* and *ech-6* in animals fed on OG1RF for 8 hours compared to OP50.

FIGURE S7

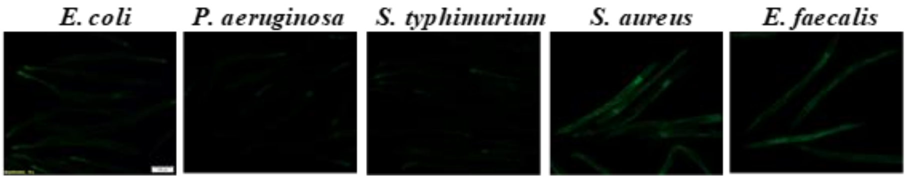


**Figure S7. Gram positive cocci diet induces *acs-2* in *C. elegans* adults.** Imaging of *acs-2P::GFP* fluorescence in animals *fed E. coli, P. aeruginosa, S. typhimurium, S. aureus* and *E. faecalis*. Scale bar, 100 μm.

FIGURE S8

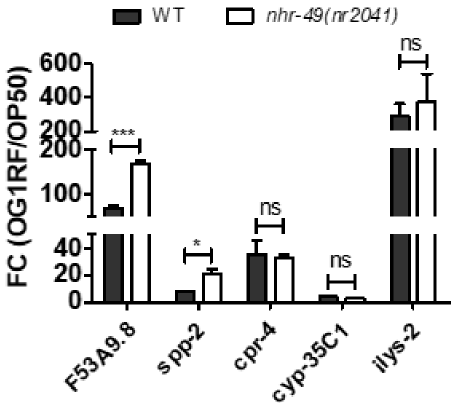


**Figure S8. Expression levels of immune effectors in *nhr-49(nr20141)* animals fed on OG1RF.** Quantitative real time PCR analysis (Mean ± SEM) of additional immune effectors in WT and *nhr-49(nr20141)* animals fed OG1RF for 8h compared to OP50.

FIGURE S9

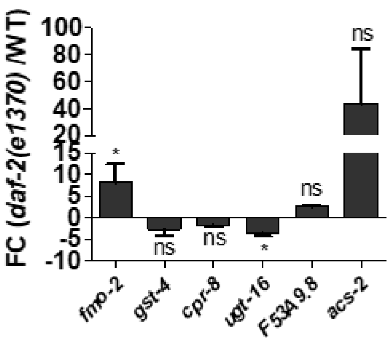


**Figure S9: Basal levels of immune effectors in *daf-2(e1370)* mutant.** Quantitative real time PCR analysis of basal levels of transcripts in *daf-2* animals compared to WT animals.
